# Reach-dependent reorientation of rotational dynamics in motor cortex

**DOI:** 10.1101/2021.09.09.459647

**Authors:** David A. Sabatini, Matthew T. Kaufman

## Abstract

During reaching, neurons in motor cortex exhibit complex, time-varying activity patterns. Though single-neuron activity correlates with movement parameters, movement correlations explain neural activity only partially. Neural responses also reflect population-level dynamics thought to generate outputs. These dynamics have previously been described as “rotational”, such that activity orbits in neural state space. Here, we find two essential features previously missed. First, the planes in which rotations occur differ for different reaches. Second, this variation in planes reflects the overall location of activity in neural state space. Our “location-dependent rotations” model fits nearly all motor cortex activity during reaching and enables higher-quality decoding of reach kinematics from single-trial spiking. Varying rotational planes allows motor cortex to more simply produce richer outputs than possible under previous models. Finally, our model links representational and dynamical ideas: a representation-like signal is present as the state space location, which dynamics then convert into time-varying command signals.

## Introduction

Controlling reaching movements requires complex, time-varying patterns of muscle activity ^1, 2^. Correspondingly, the responses of neurons in motor cortex are complex, time-varying, and heterogeneous during reaching ^2–5^. Understanding these responses and their significance for movement has been the focus of previous studies, which have found correlations between single-neuron responses and reach direction ^6, 7^, distance ^8, 9^, speed ^10^, curvature ^11^, load ^12, 13^, individual muscle activations ^14–16^, and external forces ^17, 18^. Despite these correlations and the causal link between motor cortex activity and movement, it has been challenging to find a high-fidelity relationship between the kinematics of reaching and motor cortex activity ^19, 20^.

Though single neuron activity is complex during motor control, dynamical systems analysis has revealed that population activity obeys relatively simple rules: activity of the neural population determines population activity moments later, consistent with motor cortex acting as a generator ^20–26^. During reaching, these rules have been argued to be “rotational” ^27–30^ or similar ^31–, 33^, with the spike rates of individual neurons containing low-frequency, sinusoidal modulation coordinated at the population-level. The dynamical systems approach has further organized our understanding of population activity by showing that preparatory activity sets the initial state for future movement-epoch dynamics ^19^ in dimensions orthogonal to outputs ^34–37;^ that a large trigger signal, identical across reach types, coincides with the onset of movement-epoch dynamics and may act as a ‘trigger’ for dynamics ^37–40;^ and that these dynamics act partly in “output-null” dimensions to produce command signals in “output-potent” dimensions that constitute the muscle readouts of neural activity ^23, 34^.

The dynamical systems framework provides a paradigm for understanding pattern generation in motor cortex, but despite its conceptual insights has fit only a fraction of the data. In particular, rotational dynamics fit only 20-50% of the movement-specific variance ^27^, and model variants have reached similar plateaus ^31^. Here, we demonstrate that this plateau results because a central feature of motor cortex dynamics was previously missed. Although motor cortex activity during reaching is indeed rotational, population-level rotations occupied substantially different planes in neural state space on different reaches. We also find that the center point of the rotations in neural state space varies across reaches, and that the variation in rotational planes correlates with this overall “location”. These findings enable several important advances: they allow us to account for virtually all neural variance in dorsal premotor (PMd) and primary motor cortex (M1) during reaching; describe a new class of dynamics that reconciles previously-conflicting interpretations of motor cortex; enable high-fidelity decoding from motor cortex activity with linear methods; and improve on rotational dynamics as previously understood by allowing for a much richer repertoire of motor cortical outputs.

## Results

In this work, we re-analyzed data from previous studies ^19, 27, 38^. Two monkeys, J and N, performed a “maze” variant of a delayed-reach task that evoked 72 different straight or curved reach types (“conditions”; Fig. 1a,b). Simultaneous single- and multi-unit electrophysiological recordings were made with a pair of 96-electrode Utah arrays in each monkey, implanted in PMd and in M1, yielding 79-118 units per array. We analyzed activity from each array and monkey separately. After aligning neural activity to movement onset, we trial-averaged and smoothed activity with a Gaussian kernel (s.d. = 20 ms) to estimate firing rates (Fig. 1c). Consistent with previous literature, many units displayed multiphasic activity beginning ∼150 ms prior to movement onset (Fig. 1d).

**Figure 1.**
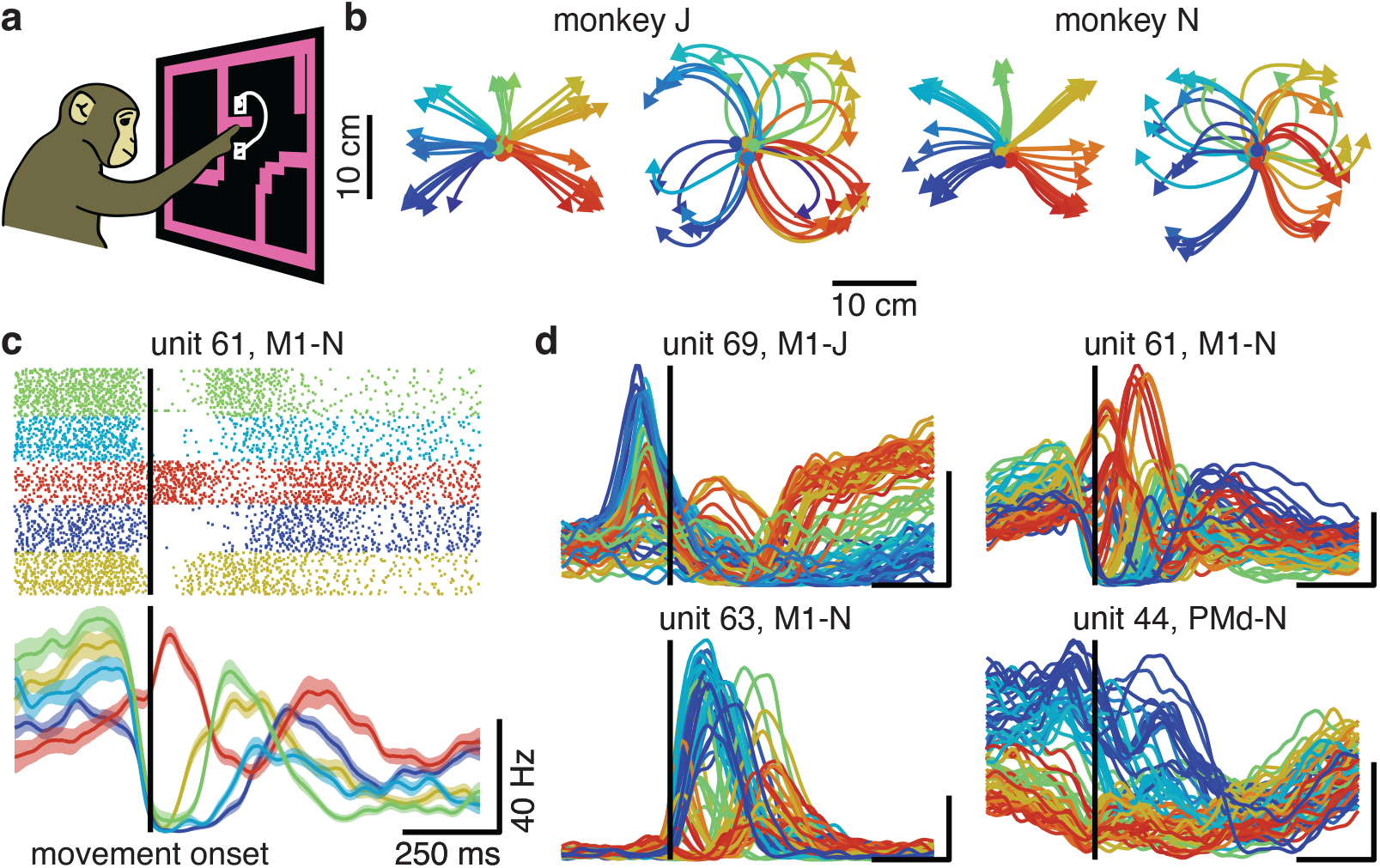
Units in motor cortex display complex activity during reaching. **a**, Illustration of task. **b**, Straight and curved reaches performed by monkey J (*left*) and N (*right*). **c**, Spiking activity and PSTH for unit 61, M1-N, for five example conditions. Lines, mean firing rates; shaded area, SEM. In PSTHs, tall vertical line indicates movement onset, and scale bars indicate 250 ms and 40 spikes/s, respectively. **d**, PSTHs for example motor cortex units, where each trace is a single condition (all conditions plotted). In **b**-**d**, colors assigned according to target angle.

### Rotational dynamics incompletely describe motor cortex activity

Previous studies have described motor cortex activity as “rotational”: containing coordinated oscillations at the level of the population ^27^. We applied the method from that work for identifying rotations in neural activity, jPCA, to our datasets to discover the planes in neural state space in which the population state rotated over time (Fig. 2a). As previously described, these planes contained coherent population-level structure, consistent with neural dynamics. As previously shown, though, these rotational dynamics explained only 7-12% (s.d. < 13% across conditions; Extended Data Fig. 1) of the variance in peri-movement firing rates (“population variance”; Methods), arguing that this model of motor cortex dynamics is incomplete.

**Figure 2:**
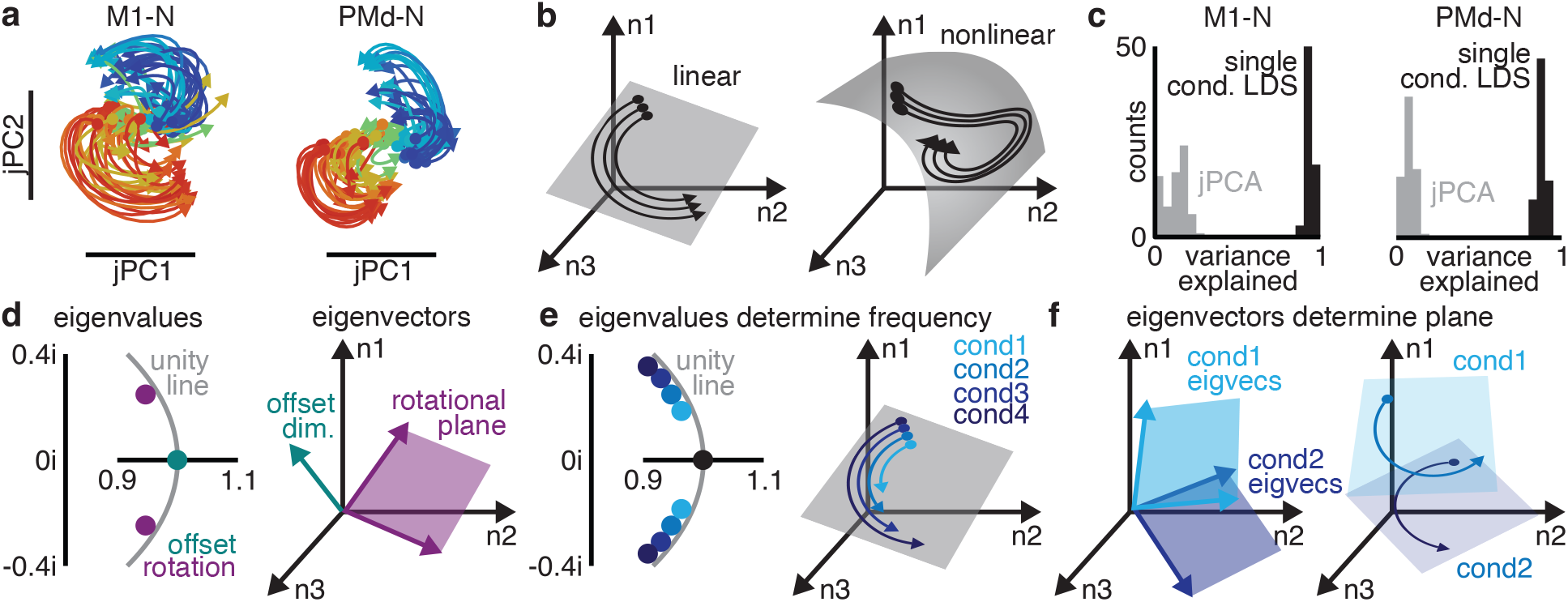
Standard rotational dynamics incompletely describe neural activity. **a**, Rotational dynamics found in motor cortex activity using jPCA, traces colored according to target angle. **b**, Examples of linear (*left*) and nonlinear (*right*) population activity. **c**, Histograms of population variance explained in motor cortex activity by rotational dynamics (jPCA, *gray*) and single condition LDSs (*black*). **d**, Linear dynamical systems can be decomposed into eigenvalues, describing rotational frequencies and half-lives (*left*); and eigenvectors, describing where the rotational planes are in neural state space. **e**, Changing eigenvalues causes rotational frequencies to change between conditions, without affecting the location of rotational planes. **f**, Changing eigenvectors causes rotational planes to differ between conditions, without affecting rotational frequencies.

We hypothesized that rotational dynamics captured only a small portion of motor cortex activity due to incorrect assumptions about motor cortex activity. jPCA finds rotational dynamics in motor cortex activity by fitting a particular class of model, a type of linear dynamical system (LDS). This restricted LDS is shared across conditions, and contains rotations at the population level. The most basic assumption of this procedure is that an LDS, rotational or otherwise, is an appropriate model for motor cortex activity. If single-condition population dynamics are highly-nonlinear or lie on a curved manifold in neural state space, an LDS will poorly describe single-condition neural activity (Fig. 2b).

As a first test of this assumption, we fit an LDS to each condition individually. This test does not make assumptions about how dynamics may be similar or different between conditions. Motor cortex activity for each single condition was low-dimensional and could be approximated as an LDS: 5-9 dimensional LDSs explained 75-93% (Fig. 2c; s.d. < 9% across conditions) of the population variance. Modeling one condition at a time with an LDS will necessarily explain more population variance than modeling them all together, and almost certainly results in overfitting. Nevertheless, the drastic gap in variance explained by rotational dynamics vs. single-condition LDSs suggests that jPCA makes some incorrect assumption about motor cortex activity.

### Rotation frequencies are conserved across reaches in motor cortex activity

If we model each condition as its own LDS, conditions might differ in two distinct ways: the frequencies of rotations and the orientations of the rotational planes (Fig. 2d). The frequency of a rotation is described by a pair of complex eigenvalues, while the rotational plane is described by the associated pair of eigenvectors. If different conditions have different eigenvalues, that means that motor cortex produces rotations of different frequencies for different reaches (Fig. 2e). If different conditions have different eigenvectors, that means that motor cortex activity rotates in different parts of neural state space on different reaches (Fig. 2f). In addition to rotations, LDSs can include an offset. A real-valued eigenvalue determines the rate-of-decay and its eigenvector determines the direction in neural state space of the offset (which we term the “state space location”). jPCA assumes both shared rotational frequencies and shared rotational planes. We therefore set out to test each assumption.

For the recorded motor cortex activity, the eigenvalues for all the conditions formed distinct clusters (Fig. 3a). These clusters corresponded to the presence of rotations on every condition with frequencies of approximately 0.5, 1.5, 2.5, and 4 Hz, along with an offset in the firing rates. The rotational frequencies (specified by the eigenvalues) were effectively identical between conditions: the variation in eigenvalues between conditions was comparable to the variation expected due to estimation noise (ROC-AUC = 0.51-0.65; Fig 3b,c) and kinematic parameters performed poorly as predictors of eigenvalues (mean R^2^ = 0.1, leave-one-out cross-validation; Extended Data Fig 2). This demonstrates that rotational frequencies are approximately conserved between conditions.

**Figure 3:**
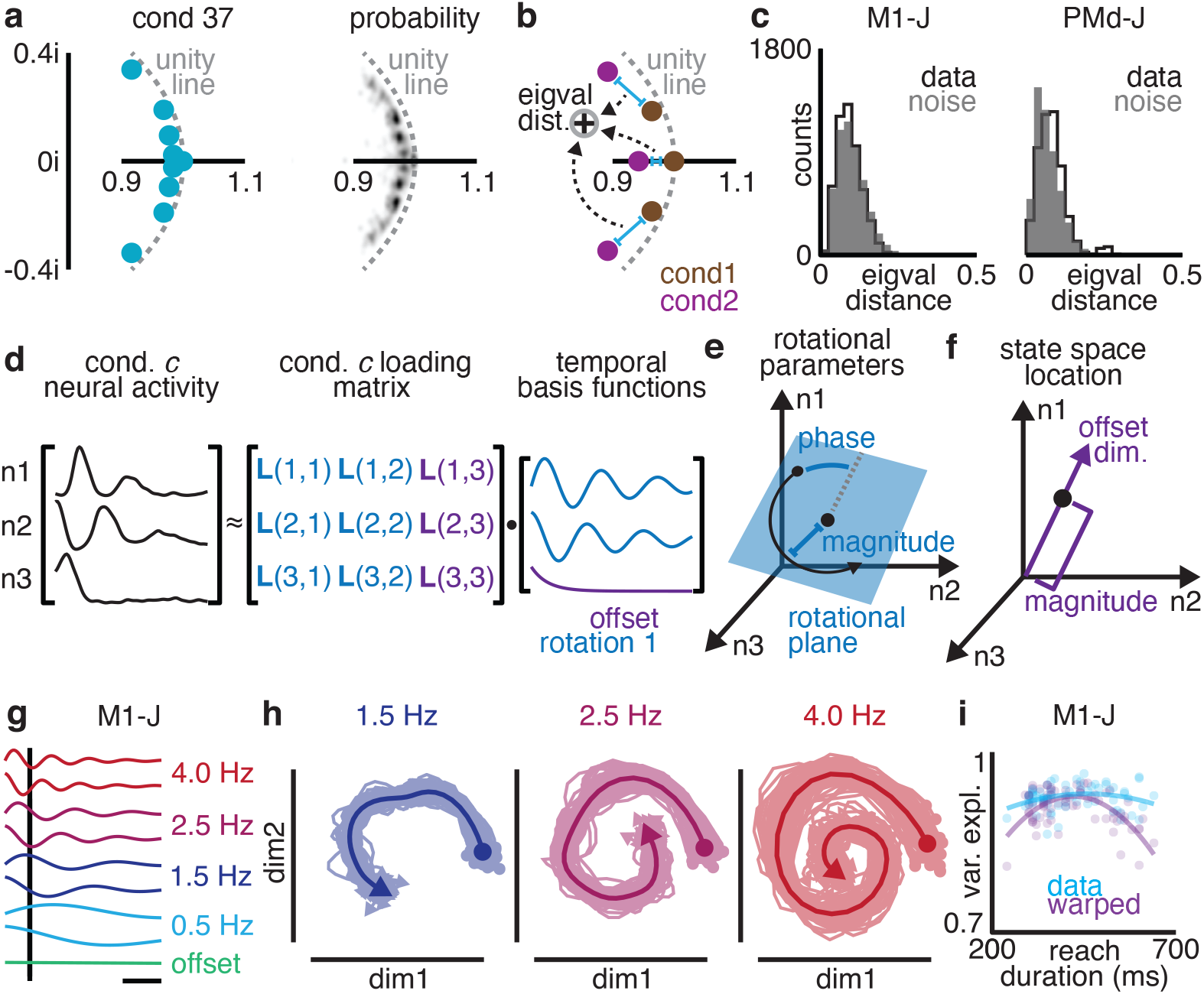
Rotational frequencies are conserved between different reaches. **a**, Eigenvalues of a single condition’s dynamics (*left*), and probability density of eigenvalues for all conditions (*right*). Data shown from M1-N. **b**, Eigenvalue distance between conditions was calculated by summing over the distances between corresponding eigenvalues in complex space. **c**, Histograms of eigenvalue distance observed between condition (*black*) and distribution expected from estimation noise (*gray*). **d**, We used a matrix factorization to identify temporal basis functions shared across conditions, along with the condition-specific loading matrix. **e**, Each condition’s loading matrix specifies the rotational plane, phase, and magnitude, or “parameters” of a rotation. **f**, Each condition’s loading matrix specifies the dimension and magnitude of firing rate offsets, or the “state space location”. **g**, Temporal basis functions identified for M1-J. **h**, Rotations recovered from motor cortex activity by projecting onto loading matrix. Solid line, mean projection across conditions; translucent lines, projections of single conditions. **i**, Warping reaches (gray) induced a negative correlation between model fit and extremity of reach duration not observed in the original data (*blue*).

### Identifying rotations with varying planes

Given that the rotational frequencies were consistent across conditions, we hypothesized that the limited fit of standard rotational dynamics was due to variation in the rotational planes between conditions. To test this hypothesis, we designed a simple method that identifies rotations in neural activity with identical frequencies for all conditions, but that allows the rotational planes and offset dimension to vary across conditions if needed. This method takes the form of a low-rank matrix factorization (Fig. 3d; Methods), in which neural activity on each condition is decomposed into two parts: a set of temporal basis functions that is identical for all conditions, and a loading matrix that can differ across conditions. Note that the factorization is performed on all conditions at once. This method fits a model of shared rotational frequencies and (potentially) varying rotational planes.

The temporal basis functions capture a limited number of patterns over time that are shared across neurons and conditions. Most of these temporal basis functions come in pairs, which form the rotations at a particular frequency, like a sine and cosine. These temporal basis functions describe rotations that are present in neural activity on every condition. The last temporal basis function is a “0 Hz” offset term. At the single-neuron level, each condition’s loading matrix describes how to recreate a neuron’s activity on that condition as a weighted sum of temporal basis functions. At the population level, pairs of columns describe the magnitude, phase, and plane of each rotation. For this reason, we refer to such pairs of columns as the parameters of a rotation (Fig. 3e). One last column of the loading matrix defines the dimension and magnitude of the offset, and so determines the condition *c*’s state space location: the point in neural state space around which the population state rotates on that condition (Fig. 3f).

This method captures motor cortex activity well. In agreement with our fits to single conditions, the optimal temporal basis functions contained 3-4 rotations at 0.5, 1.5, 2.5, and 4 Hz, along with an offset (Fig. 3g; optimal number determined by cross-validation). These rotations explained 90-96% of the population variance, much greater than expected by chance (s.d. < 2% across conditions; smoothed temporal shuffle, 35-46%, see Methods; Wilcoxon Rank Sum Test, p < 0.05).

Two potential degeneracies in this model must be controlled for. First, one concern is that this factorization may act as a kind of special-purpose Fourier transform: not identifying true rotations in motor cortex activity, but simply performing a frequency decomposition. In this case, we would expect any given plane in neural state space to contain a mixture of different frequencies, and therefore each rotation frequency would not be cleanly “recoverable” from its plane of neural activity (Methods). Instead, as expected from the model, motor cortex activity on each condition was cleanly rotational at the expected frequencies in the expected planes of neural state space (Fig. 3h). The projections from each condition’s motor cortex activity recovered the temporal basis functions almost perfectly (91-95% variance explained; s.d. < 2.5% across conditions; chance, 63-80%; Wilcoxon Rank Sum Test, p < 0.001), arguing the factorization in fact identified rotations in motor cortex activity.

The second concern is that a common time course could be present in motor cortex activity simply because different reaches take similar amounts of time. If similar reach time courses were the primary source of the similar neural frequencies across conditions, then warping reaches to identical durations should further improve this similarity. Prior to warping, the frequencies of the rotations were unrelated or weakly related to reach duration (M1-N, Pearson’s Rho = −0.26, p = 0.023; other datasets, Pearson’s Rho = −0.18-0.21, p > 0.067). Warping to equalize reach duration induced a negative correlation: the more a reach was warped, the less that condition’s neural activity was fit by the common rotations (Fig. 3i; Pearson’s Rho = −0.63 to −0.52, p < 0.001). The conserved rotational frequencies were therefore not explained by the similar time courses of the reaches. Given that the fit of this model could not be explained by degeneracies, we used the rotational planes identified by it for subsequent analyses.

### Rotational planes differed across reaches

To quantify how different the rotational planes were on different conditions, we first used the alignment index ^35, 41^. The alignment index measures how well one subspace in neural state space (such as a plane) aligns with another, measuring 1 if the subspaces are identical, 0 if the subspaces are completely orthogonal, or intermediate values for circumstances in between (Fig. 4a). We compared the corresponding rotational planes for pairs of conditions: for example, we calculated the alignment index between the 4 Hz rotational planes on two different conditions. We found that rotational planes were far less aligned than expected due to estimation noise (ROC-AUC = 0.85-0.96; Wilcoxon Signed Rank Test, p < 0.001; Fig. 4b; Extended Data Fig. 3a). In other words, the orientation of the rotation planes differed across condition, contrary to the assumptions of previous studies on rotational dynamics in motor cortex.

**Figure 4:**
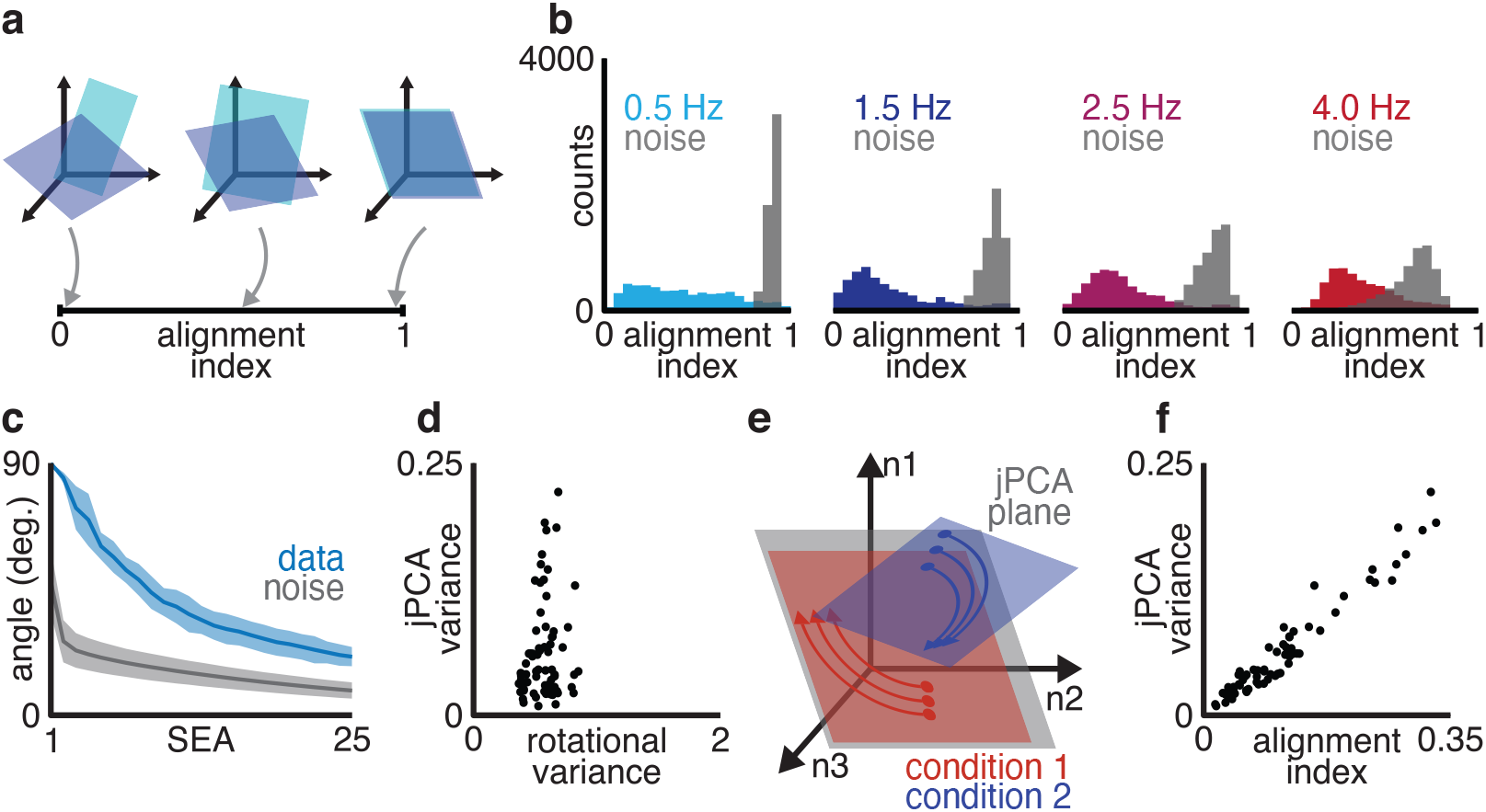
Rotational plane differs substantially between reach conditions. **a**, The alignment index quantifies the overlap between two rotational planes. **b**, Alignment indices between corresponding (same-frequency) rotational planes on pairs of reach conditions (*colored*) and distribution expected by estimation noise (*gray*). **c**, Subspace excursion angles for corresponding rotational planes across conditions (*blue*), along with angles expected by estimation noise. Line, mean; shaded, 1 standard deviation. **d**, Total rotational variance of a condition vs. the condition’s variance in the first jPCA plane. **e**, The “misalignment” hypothesis: all conditions are equally rotational, but some rotate in planes orthogonal to the jPCA plane. **f**, Alignment index between a condition’s rotational planes and the first jPCA plane, vs. the condition’s variance captured by jPCA. Data shown from M1-J.

To better understand how much each rotational plane varied, we used a novel metric, Subspace Excursion Angles (SEA). SEA takes a set of subspaces in neural state space (for example: the 4 Hz rotational plane of each condition) and sorts the set so that each subspace in the list has the largest possible subspace angle with respect to all the previous subspaces in the list. SEA therefore quantifies how many dimensions contain substantial variation in subspace. Across conditions for each rotational frequency, SEA identified 6-20 distinct dimensions with angles > 45° to each other in motor cortex activity (Fig. 3c; Extended Data Fig. 3b). Rotational planes therefore varied substantially between conditions.

These results explain a previous observation: while most conditions exhibit rotations in the jPCA-identified rotational planes, on many conditions neural activity has very low magnitude in the rotational plane. We found that these conditions were not actually less “rotational”: variance in the jPCA planes was only moderately correlated with the variance in the rotations we identified (Fig. 4d; Pearson’s Rho = 0.39-0.46, 66% of jPCA planes significantly correlated at p < 0.05). Instead, for conditions that did not exhibit clear rotations in the jPCA planes, the rotations our model discovered were misaligned with the jPCA planes (Fig 4e,f; Pearson’s Rho = 0.82-0.97 between plane alignment and variance explained by jPCA, p < 0.001). Previously described rotational dynamics were therefore an incomplete description of motor cortex activity: while jPCA found planes in which motor cortex activity rotated for some reach conditions, these planes were compromises that missed equal-magnitude but nearly-orthogonal rotations on other conditions.

### Different-frequency rotations occupied distinct yet coordinated subspaces

Although the rotational planes varied substantially between conditions as shown above, we found that this variation was structured to avoid reusing the same plane for different frequency rotations. If we considered pairs of conditions and computed the alignment index for the corresponding rotations (e.g., the 2.5 Hz plane for both conditions) vs. the alignment index for different-frequency rotations (e.g., 2.5 Hz for one condition and 4 Hz for the other), the different-frequency pairs were consistently nearly-orthogonal, and the corresponding rotations were better aligned (Fig. 5a; Wilcoxon Signed Rank Test, p < 0.001; Extended Data Fig. 4). Different frequencies were therefore segregated into different subspaces of neural state space.

**Figure 5:**
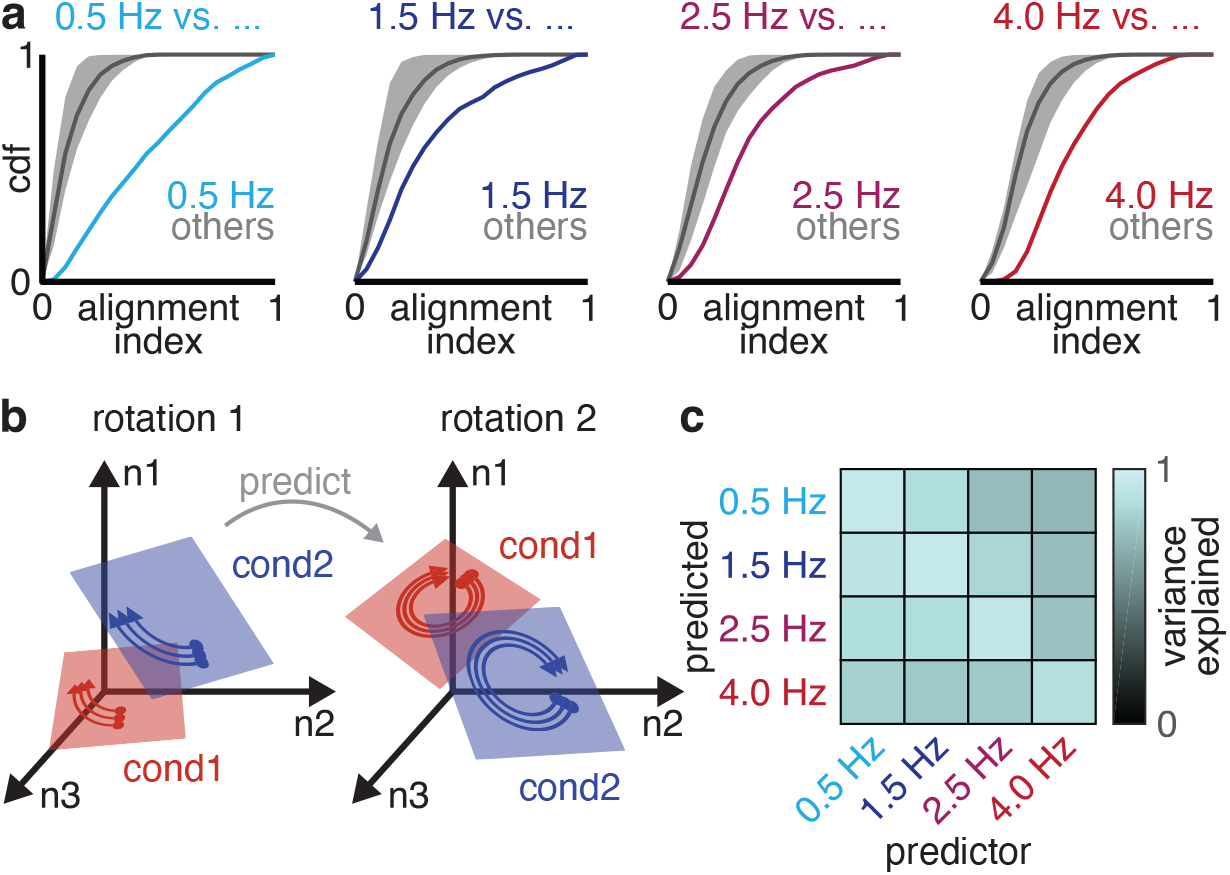
Rotational planes coordinated variation between conditions. **a**, Cumulative distribution function of alignment index between different-frequency rotational planes (*gray*) and same-frequency rotational planes (*colored*) on pairs of conditions. Same-frequency rotational planes were more aligned than different-frequency rotational planes. **b**, To test for correlations between different rotations’ parameters, we used one rotation’s parameters to predict a different rotation’s parameters. **c**, Mean variance explained in each rotation across condition by prediction from a separate rotation. Data shown from M1-J.

Although different frequency rotations occupied different subspaces of neural state space, their orientations across conditions were nevertheless related. We tested whether one rotation’s parameters for a given condition could be used to predict the parameters of a different-frequency rotation in the same condition. For example, could the 4 Hz rotation’s parameters be predicted from the 2.5 Hz rotation’s parameters using the same model for all conditions (Fig. 5b)? We quantified the prediction quality as the variance explained in the true rotation by the predicted rotation, using the same prediction model for all conditions. We quantified this predictive ability over all pairs of rotation. Each rotation predicted an average of 43-94% of the variance in other rotations (s.d. < 25%, 6-fold cross validation), significantly greater than expected by chance (Fig. 5c; Wilcoxon Signed Rank Test, p < 0.001). This finding suggests that although rotations occupied separate parts of neural state space, their parameters are set jointly by an underlying low-dimensional structure.

### Motor cortex activity features location-dependent rotations

So far, we have analyzed differences in the orientations of the rotational planes between conditions. Our results generalized to the state space location, the point around which the rotations occur. The state space location subspace was multi-dimensional: the dimensions occupied across conditions were less aligned than expected due to estimation noise (ROC-AUC = 0.88-0.95; Wilcoxon Signed Rank Test, p < 0.001; Extended Data Fig. 5a). To determine the dimensionality of the subspace, we again used SEA. SEA identified 4-5 state space location dimensions with angles > 15° (Extended Data Fig. 5b). Moreover, the state space location dimensions were distinct from those of rotations (same analysis as Fig. 5a; Wilcoxon Signed Rank Test, p < 0.001; Extended Data Fig. 5c).

In a dynamical system, location in state space determines dynamics. Accordingly, the state space location was strongly predictive of rotations’ parameters (Fig. 6a). We repeated our previous analysis, but instead used the state space locations to linearly predict rotations’ parameters. The state space locations predicted an average of 67-80% of the total population variance (Fig. 6b; s.d. < 15%, 6-fold cross validation), significantly greater than expected by chance (Wilcoxon Signed Rank Test, p < 0.001). These findings suggest that motor cortex activity uses “location dependent rotations” (LDR): that “where” the population state is in the neural state space determines “how” the population state rotates over time – that is, in which dimensions the rotations occur.

**Figure 6:**
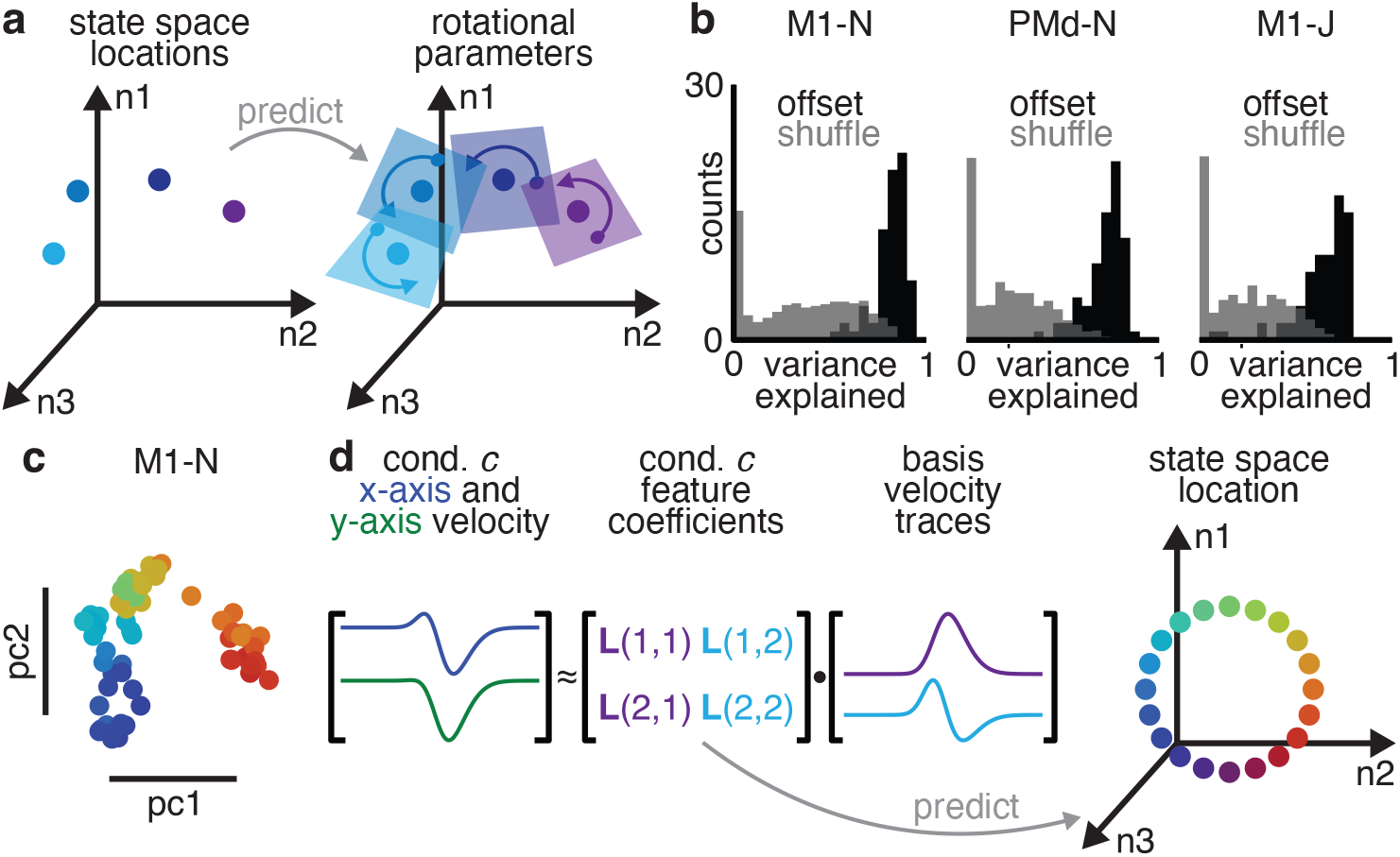
State space location predicts rotational plane. **a**, Illustration of “location-dependent rotations” model. The state space location was used to predict the parameters of every rotation. **b**, Population variance explained by predicting rotational parameters from state space location. **c**, PCA plot of state space locations across condition. Each dot is a single condition. **d**, Reach parameters describing reach angle and curvature were used to predict state space location.

### Reach kinematics correlate with state space location

If motor cortex does in fact use location dependent rotations, then the state space location determines the time-varying aspects of neural activity. In agreement with the many known relationships between motor cortex activity and movement parameters, the state space location was strongly related to reach kinematics. We visualized state space locations across conditions using Principal Component Analysis (PCA). Qualitatively, state space location related to reach angle in a continuous manner (Fig. 6c). To quantify the relationship between state space location and reach kinematics, we used linear regression to predict state space location from kinematics (Fig. 6d). We used linear dimensionality reduction to summarize reach kinematics, extracting four “reach parameters” per condition that captured information about reach angle and curvature over the course of the reach. Reach parameters linearly predicted 37-71% of the variance in state space location (6-fold cross validation), greater than expected by chance (shuffle, p < 0.001). Allowing nonlinearities via kernel regression raised the variance explained to 69-81% of the variance in state space (6-fold cross validation; shuffle, p < 0.001). These observations suggest that the state space location relates closely to reach kinematics.

### Reach kinematics correlate with the dynamics of motor cortex activity

Above, we found that the state space location of motor cortex activity could predict the parameters of rotations in neural activity, and that state space location relates tightly to reach kinematics. We therefore hypothesized that motor cortex activity during reaching could be predicted by using reach parameters (a whole-sequence summary of kinematics) to reconstruct both the state space location and the rotational features (orientation, magnitude, and phase for each frequency) of the neural activity. Most previous attempts to relate neural activity to movement have used moment-by-moment correlations between kinematic parameters and firing rates to fit an instantaneous encoding model. These approaches have explained at most approximately half the variance of motor cortex activity ^19, 42^, leading to discussion of what the rest of the activity in motor cortex corresponds to ^20^. Our proposed LDR encoding model takes a different approach, predicting features of motor cortex dynamics (the state space location and rotational parameters) instead of activity directly. Moment-by-moment firing rates are determined only indirectly, by the rotations unfurling over time in the specified planes.

To fit this model, we used linear regression to predict the loading matrix of a condition’s neural activity, which includes both the state space location and rotation parameters, from that condition’s reach parameters (which summarize the kinematics; Fig. 7a). Importantly, a single model was used for all conditions. Motor cortex activity was reconstructed for held-out conditions by first predicting the loading matrix, then using the temporal basis functions to produce the predicted neural activity over time. We quantified model fit as the population variance explained in motor cortex activity.

**Figure 7:**
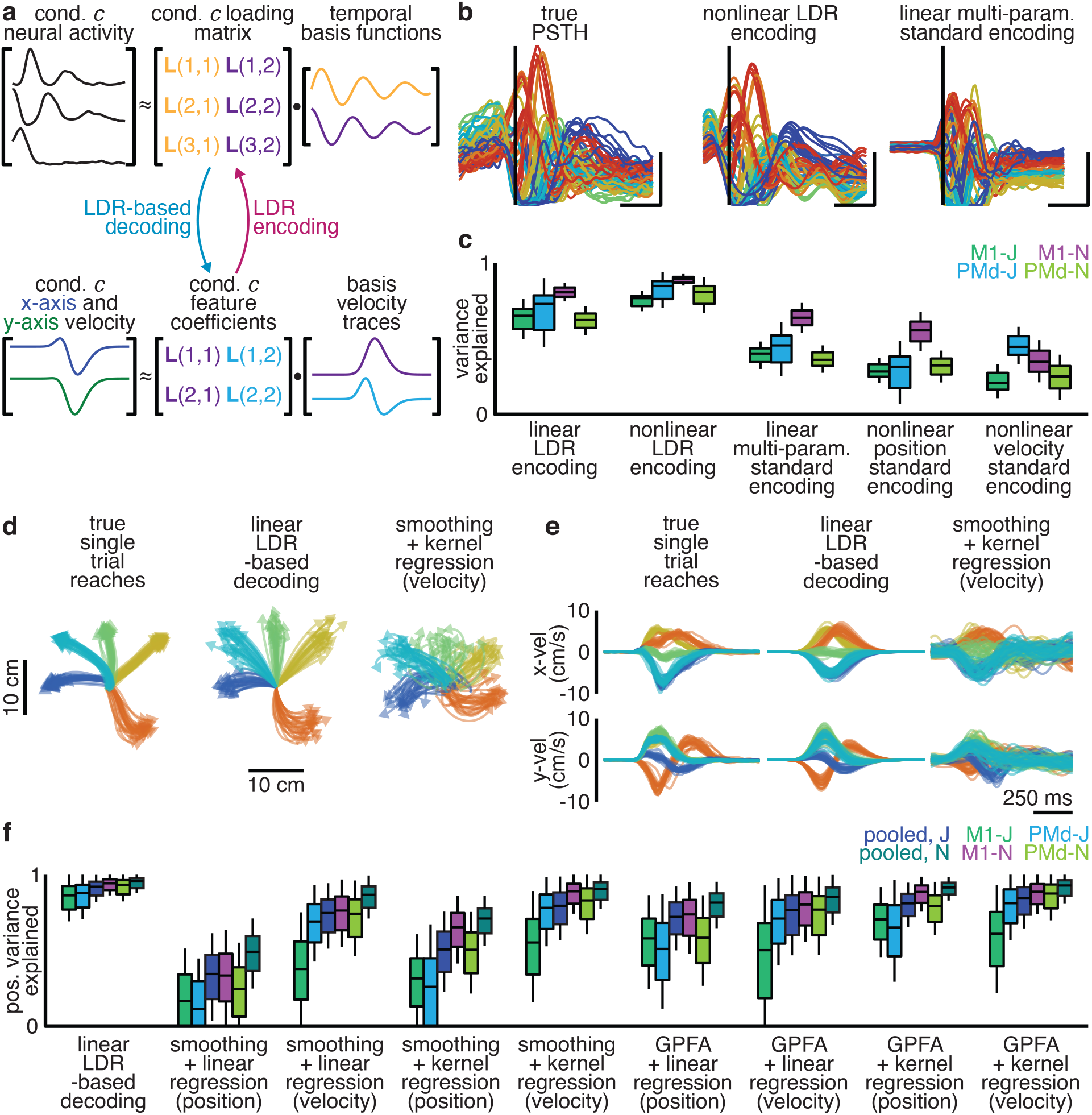
Reach kinematics are encoded in the rotational planes and state space location of motor cortex activity. **a**, Illustration of LDR encoding and decoding model. **b**, True PSTH for unit 61, M1-N (*left*), reconstruction by a nonlinear LDR-encoding model (*center*), and reconstruction by a standard linear multi-parameter standard encoding model (*right*). **c**, Median (bar), interquartile range (IQR, box), and 1.5x IQR (whiskers) of population variance explained by linear and nonlinear LDR-encoding models, along with variance explained by standard encoding models. **d,e**, Hand position over time (d) and hand velocity over time (e) along with reconstruction by LDR-based decoding and kernel regression on smoothed spiking (100 ms Gaussian filter). **f**, Median, IQR, and 1.5Ix QR of variance in hand position explained by LDR-based decoding, along with variance explained by standard techniques employing a de-noising step and then fitting an instantaneous activity-based decoder of reach kinematics. LDR-based decoding explained more variance than every other method tested (p < 0.001).

This linear LDR-encoding model explained most of the population variance: 60-78% (s.d.< 22%, 6-fold cross-validation), greater than expected by chance (Wilcoxon Signed Rank Test, p < 0.001). To determine whether including a nonlinearity could further boost performance, we also tested kernel regression. This nonlinear LDR encoding model explained 71-87% of the population variance (s.d. < 17%, 6-fold cross-validation), greater than expected by chance or memorization of the dataset (Fig. 7b,c; Wilcoxon Signed Rank Test, p < 0.01; Extended Data Fig. 6). Motor cortex activity could therefore be predicted from kinematics by predicting the state space location and rotations’ parameters.

To reiterate, LDR encoding models do not fit an instantaneous relationship between neural activity and kinematics as in standard models. LDR predicts the state space location and rotations’ features from a summary of the kinematics for the entire reach, and thus forms a “sequence-to-sequence” encoder: an entire reach is used to predict features of motor cortex dynamics. This difference is crucial. We quantified the performance of various linear and nonlinear standard encoding models and found they were substantially outperformed by both linear and nonlinear LDR encoding models (Fig. 7b,c; Wilcoxon Signed Rank Test, p < 0.001). Importantly, though, this model is compatible with motor cortex having a subset of dimensions that relate instantaneously to muscle activity or kinematics (see Discussion).

### LDR allows linear decoding of single-trial kinematics

We hypothesized that because LDR relates a larger fraction of neural activity to kinematics, it would also allow us to better decode reach kinematics from neural activity. We therefore formulated an LDR-based decoder to predict reach kinematics from single-trial spiking activity, which is essentially the encoding model in reverse applied to single-trial spiking data (Fig. 7a, Methods). In particular, it acts as a linear sequence-to-sequence decoder: projecting spiking activity onto the temporal basis functions to extract the state space location and rotational dynamics orientations, and using these to predict a static representation of the reach trajectory.

Despite using only linear operations on spiking activity, this LDR-based decoder on average explained 84-92% of the variance in the hand’s position over time (s.d. < 15% across trials; 2-fold cross validation across conditions). This decoder outperformed every other decoder we tested, including combining advanced de-noising methods with nonlinear decoders (Fig. 7d-f; Wilcoxon Signed Rank Test, p < 0.001 vs. every other tested decoder for all datasets).

Because the LDR-based decoder was trained using trial-averaged data, one concern might be that LDR-based decoding may produce only the average reach for each condition – de-noising trajectories at the expense of real variability. Using an LDR-based decoder, however, we were able to decode trial-by-trial variations in reach curvature (Pearson’s rho, 0.16-0.41; p < 0.001), indicating that our model decoded fine variations in movement.

## Discussion

Our results reveal several previously-undescribed features of motor cortex activity in the monkey during a variety of reaching movements. First, although the same frequencies of rotations were present across reaches, the rotational planes changed substantially across different reach conditions. Second, the overall state space location of motor cortex activity predicts which planes are occupied by the rotations in the population state. Finally, both the state space location and the rotational planes correlate strongly with the kinematics of the ongoing reaches.

Our finding that rotational planes differ between different reaches explains why previous rotational models of motor cortex failed to describe the activity for many conditions: it is not that dynamics were weak for those conditions, but simply that rotations occurred in other dimensions of neural state space. This variability in rotational plane means that neural dynamics were not, as previously assumed, very low-dimensional. This is consistent with an emerging understanding that motor cortex activity is higher-dimensional than previously hypothesized ^16^.

Consistent with our observations, several recent studies have documented rotational planes varying between conditions. Motor cortex has been observed to rotate in largely separated planes during forward and backward cycling ^29^, while network models of motor cortex exhibit different rotational planes during different grasping movements ^43^. These observations fit with a growing body of literature suggesting that confining different dynamics to different parts of state space allows recurrent neural networks to flexibly perform multiple behaviors without catastrophic forgetting ^44^, to reduce interference between different movements ^45^, or to segregate movement-epoch dynamics from preparatory activity ^38^.

We observed that rotational planes not only differ between conditions, but the differences vary smoothly as a function of difference in reach kinematics. This reconciles seemingly-contradictory observations about the dimensionality of neural data ^22^. The variability in rotational planes makes motor cortex activity moderately high-dimensional across neurons. Yet at the same time, that rotational planes change smoothly as function of reach kinematics means that motor cortex activity is low-dimensional in conditions, because there are a limited number of independent neural responses motor cortex can produce ^22, 46, 47^.

This low-dimensionality of conditions, combined with high neural dimensionality due to rotational plane variation, may allow motor cortex to generalize well while nevertheless being sufficiently expressive ^48, 49^. Smoothly varying the rotational plane may allow motor cortex to generate the “correct” activity patterns for new reaches by reaping the benefit of strong generalization due to local linearity ^50, 51^. On the other hand, allowing the population state to rotate in substantially different planes for different reaches may allow motor cortex to produce a variety of different output signals across reaches ^33^. Qualitatively, the ability of dynamics to change systematically with state space location also relate to work on network solutions to producing a variety of cycling speeds, where speed is encoded along a dimension in the network’s state space, while dynamics vary along the encoding dimension ^52^; and to how the brain times intervals ^48^.

Our findings additionally help reconcile classic representational models of motor cortex with the dynamical systems perspective. Representational models focus on firing rates correlating with movement parameters such as the direction and distance of a target. In our model, this tuning is captured in the correlation between location in state space and parameters of movement. Our findings suggest the state space location is used to set up motor cortex dynamics to produce the appropriate trajectory in neural state space, thereby converting a simple representation into the necessary time-varying outputs. This model therefore argues that tuning and dynamics are meaningful and interrelated.

Based on our findings, we propose the following conceptual model of motor cortex during reaching (Fig. 8a). The population state of motor cortex is moved to the appropriate state space location, and the population state then rotates in planes set by the state space location. The phase and magnitude of the rotation are set by the initial activity, whether this is from a preparatory period or the irreducible preparation preceding movement ^19, 35, 37^. We speculate that these rotations are generated by autonomous dynamics (though see ^23^ for caveats), so that the local flow field of motor cortex at each state space location determines the rotational planes. As in many models, the outputs from cortex in our model are determined by the activity in specific output-potent dimensions ^34^, which may contain only modest variance ^16, 18^. We speculate that having the system work in this way may help the brain solve the “inverse problem” ^53^: it allows a linear mapping of the desired kinematic trajectory to a location in neural state space, and the local dynamics can then produce the needed patterns in the output dimensions.

**Figure 8:**
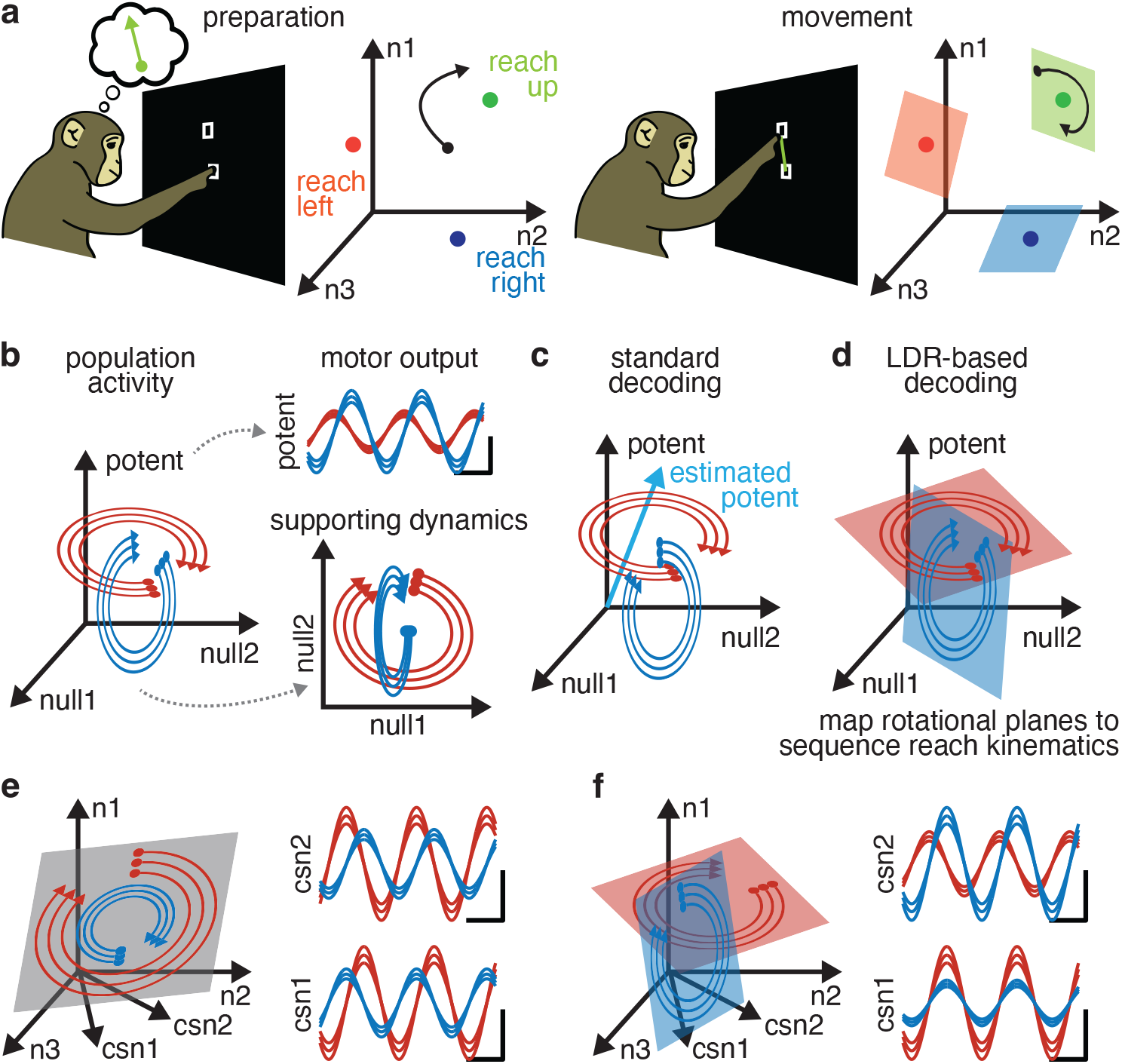
LDR allows richer outputs than rotational dynamics. **a**, Illustration of proposed model. Preparatory activity moves the population state to the necessary state space location, where local rotational dynamics cause the population state to rotate in the correct planes upon movement onset. **b**, In a dynamical system, output-null dimensions act as supporting dynamics for producing activity along output-potent dimensions. This means activity in output-null dimensions, though not directly “read out” by the nervous system in driving behavior, is informative of behavior. **c**, Standard decoders attempt to estimate output-potent dimensions, but in the process discard information in output-null dimensions. **d**, LDR-based decoding, by using motor cortex dynamics on each condition, leverages output-null and output-potent activity to refine estimates of reach kinematics. **e**, Rotational dynamics impose strict phase- and magnitude-locking on outputs, such as CSNs. **f**, By contrast, LDR allows for CSNs to break phase- and magnitude-locking, allowing motor cortex to produce a richer set of outputs.

This conceptual model explains why an LDR encoding model is able to explain more variance in neural activity than standard encoding/representational models. Dynamical systems models argue that neural activity occupies both output-potent dimensions, which directly drive outputs, and output-null dimensions, which are needed to support the generating dynamics (Fig. 8b) ^23, 25, 34^. Standard encoding models consider correlations between neural activity and movement variables, and therefore can only explain activity in output-potent dimensions. LDR encoding models relate whole neural trajectories to whole-reach kinematics, and are therefore able to predict activity in both output-potent and output-null dimensions from a high-level description of the reach. Similar logic applies to decoding models. Though standard decoders can use output-potent dimensions to decode movement from neural activity, LDR-based decoding can leverage output-null activity as well, effectively giving it more data to work with (Fig. 8c,d) ^54^. To do so, however, requires the additional understanding provided by LDR of the relationship between representation and dynamical pattern generation.

LDR may have other advantages for the brain, allowing motor cortex to generate richer command signals during movement than would be possible with strictly rotational dynamics. When limited to a single readout, rotational dynamics can approximate any arbitrary pattern over time given a well-chosen initial state. This allows rotational dynamics to drive the needed spiking in, for example, a single corticospinal neuron (CSN). Rotational dynamics, however, cannot arbitrarily set the phases and amplitudes of two or more CSNs’ activity. Readouts from rotational dynamics are strictly locked in phase and magnitude, meaning that spiking activity of multiple CSNs cannot be independently modulated (Fig. 8e). LDR does not have this limitation. With rotations oriented appropriately in state space, each CSN contains oscillations of the correct amplitude and phase to produce the needed pattern of spiking over time (Fig. 8f). By changing rotational planes between conditions, CSNs can be driven with effectively independent phases and magnitudes.

Our findings here suggest several immediate avenues of future research. LDR-based decoding, as a sequence-to-sequence model, cannot be used for real-time control of brain computer interfaces (BCI). Our findings suggest, however, that using motor cortical dynamics in both output-potent and -null spaces to decode movement condition may be more effective and robust than decoding instantaneous kinematics directly. Our approach in this paper may additionally be useful for studying strongly “tangled” systems ^29^ that have resisted dynamical systems analysis ^55^. Finally, by providing a link between neural state location, dynamics, and behavior, LDR may provide insight into how difficult transformations between inputs and outputs occur in real and artificial neural networks.

## Acknowledgements

This work was supported by funding from The University of Chicago, the Neuroscience Institute at The University of Chicago, the Alfred P. Sloan Foundation, and NIH R01 NS125270-01 (MK). We thank C. Pandarinath for suggesting controls, and S. Bensmaia and J. MacLean for helpful feedback on the manuscript.

## Methods

### Subjects, behavior and data pre-processing

All data used here have been analyzed in previous studies. Animal protocols were approved by the Stanford University Animal Care and Use Committee. Two adult male rhesus macaque monkeys (*Macaca mulatta*), J and N, performed the “maze” center-out delayed-reaching task for juice reward ^19^. Briefly, the monkey initially fixated a central point on a vertical screen with the eye and a cursor floating slightly above the fingertips, then a slightly jittering target was shown. After a brief delay (0-1000 ms), a Go cue was presented (cessation of target jitter, filling in of target, and disappearance of fixation point), and the monkey was required to make a fast, accurate reach to the target. On some trials, the monkey was required to avoid virtual barriers presented at the same time as the target, eliciting curved reaches to produce 72 different reaching “conditions” (36 straight, 36 curved). For both monkeys, data were collected using two 96-electrode “Utah” arrays (Blackrock Microsystems, Salt Lake City, UT), one implanted in PMd and one in M1. Both single-units and stable multi-units were included for analysis. Spiking activity was binned at 1 millisecond, smoothed with a 20 ms Gaussian kernel, trial-averaged, and down-sampled to 10 millisecond resolution. Except where noted, all analyses were performed on each dataset (monkey and array) independently.

### Fitting dynamics to a single condition

Condition-specific dynamics were examined by fitting, for each condition independently, a discrete-time linear dynamical system to population activity from 150 ms prior to movement onset to 850 ms after movement onset. If an eigenvalue in a fit LDS had a half-life >10 seconds, it was reduced to 10 seconds. To lessen overfitting, we first reduced the dimensionality of single-condition neural activity by projecting activity on to the top *k* singular vectors in neural state space. We determined the number of singular vectors to use via cross-validation (see below; ^56^). To quantify the fit of linear dynamics, for each condition we simulated the dynamics forward from the population state 150 ms prior to movement onset, then quantified the population variance explained in the observed neural activity for that condition.

To perform the cross-validation mentioned above, for each condition independently we assigned trials into one of two partitions, then smoothed and trial-averaged each partition independently to produce independent estimates of firing rates. We denote these estimates **X** and **Y**. We estimated the singular vectors of population activity from **X**. We then, for different dimensionalities *k*, projected **X** on to its top *k* singular vectors and quantified the variance explained in **Y**. We repeated this procedure 100 times per condition, and selected the value of *k* that maximized the average variance explained across folds and conditions.

### Controlling for the expected similarity of conditions’ eigenvalues

There are two reasons that eigenvalues for pairs of conditions may differ: true differences and estimation noise. To quantify the expected dissimilarity due to estimation noise, we simulated 2 sets of 20 trials from each condition’s firing rate via a Poisson process, then smoothed and trial-averaged each partition to estimate firing rates. We then fit linear dynamics independently to each partition. Finally, we quantified the distance between the eigenvalues between the two estimates of the same condition’s firing rates. This provides an estimate of the eigenvalue distance attributable to estimation noise under the Poisson assumption. We repeated this procedure 500 times per condition to estimate a distribution of eigenvalues distance. We then compared this null distribution to the true distribution of eigenvalue-distances between pairs of conditions. We quantified differences in distribution using a Wilcoxon Rank-Sum Test to test for differences in medians of the distribution, along with quantifying the Receiver Operator Characteristic Area Under the Curve (ROC-AUC) to quantify the discriminability between the two distributions.

### Controlling for the expected similarity between conditions’ eigenvectors

The alignment index provides a measure of similarity between the eigenvectors of two separate conditions. There are two reasons that two conditions’ eigenvectors may have a low alignment index: true differences and estimation noise. To quantify the expected alignment index due to noise, as above we simulated 2 sets of 20 trials from each condition’s firing rate via a Poisson process, then smoothed and trial-averaged each partition to estimate firing rates. We then identified the top *k* neural singular vectors of each partition, where *k* is the earlier identified dimensionality, and quantified the alignment indices between the two partitions. We used the singular vectors, as the eigenvectors and singular vectors are equivalent up to a linear transformation, and the alignment index is invariant to linear transformations. We repeated this procedure 500 times to estimate a distribution of alignment indices purely from single-trial noise. We then compared this null distribution to the true distribution of eigenvector alignment indices between pairs of conditions. We quantified differences in distribution using a Wilcoxon Rank-Sum Test to test for differences in medians of the distribution, along with quantifying the Receiver Operator Characteristic Area Under the Curve (ROC-AUC) to quantify the discriminability between the two distributions.

### Factorizing neural activity using temporal basis functions

To identify rotations conserved in motor cortex activity across conditions, without assuming preservation of rotational planes across conditions, we exploited the convenient fact that any diagonalizable LDS,

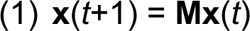

can be re-written on a single condition (defined by a single initial state) as,

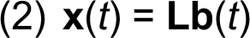

expressed as the product of loading matrix **L** and a set of temporal basis functions **b** (see Supplement for derivation). Briefly, the loading matrix captures information about the eigenvectors and initial state of the LDS, while the temporal basis functions describe the temporal patterns produced by the eigenvalues. Importantly, two LDSs with the same eigenvalues will have the same temporal basis functions, even if their eigenvectors differ. Assuming neural activity on each condition has the same eigenvalues, but different eigenvectors, these temporal basis functions will be shared by neural activity across conditions, just with (potentially) different loading matrices. Neural activity on each condition is therefore just a (potentially) different linear transformation on the same temporal basis functions, so the temporal basis functions can be recovered via a Singular Value Decomposition on the neural activity across conditions concatenated into a (*neurons* x *conditions*)-by-*time* matrix. The top *k* singular vectors over the *time* index are assigned as the temporal basis functions, while the top k singular vectors over *neurons* and *conditions* (weighted by the singular values) can be partitioned into condition-specific loading matrices. Activity on condition *c*, a *neurons*-by-*time* matrix **X**(c), is then expressed as

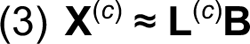

where **L**(c) is a *neurons*-by-*k* loading matrix for condition *c*, and **B** is a *k*-by-*time* matrix where each row of the matrix is a temporal basis function. This procedure is related to the Higher-Order Singular Value Decomposition. To quantify the approximation provided by this factorization, for each condition we multiplied that condition’s loading matrix by the shared temporal basis functions and computed the population variance explained in that condition’s neural activity.

### De-mixing temporal basis functions

Like many matrix factorizations, the factorization in equation 3 is only unique up to an invertible linear transformation. To allow for later analyses that considered the different frequencies of rotations, we aimed to “de-mix” the temporal basis functions into as pure of frequencies as possible – corresponding to functions attributable to single (pairs of) eigenvalue(s). To this end, we fit an LDS to the temporal basis functions from 150 ms prior to movement to 850 ms after movement onset and projected the temporal basis functions onto the eigenvectors of this LDS,

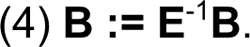

This operation does not affect the quality of the factorization, as we correspondingly multiplied the loading matrix of each condition by the eigenvectors,

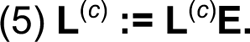

The eigenvalues of the fit LDS can then be matched one-to-one with temporal basis functions.

### Cross-validating the number of temporal basis functions

To cross-validate the number of temporal basis functions to use, we assigned trials to one of two partitions, then smoothed and trial-averaged each partition to produce independent estimates of each condition’s firing rates. We estimated loading matrices and temporal basis functions for one partition, varying the number of temporal basis functions used. We then reconstructed, for each condition, the neural activity as the product of that condition’s loading matrix and temporal basis functions. We repeated this procedure over 100 folds, and selected the number of temporal basis functions to maximize the average variance explained in the held-out partition across conditions. After the fact, we visually screened temporal basis functions, and discarded the last basis function if it was “unpaired”; that is, if it did not have a similar-frequency basis function with a phase offset.

### Quantifying the recoverability of temporal basis functions

Basis function decompositions, such as the Fourier transform, have been applied widely to decompose complicated datasets into linear sums of more interpretable functions. We therefore needed a method to ensure that the factorization described above was identifying rotations in motor cortex activity, and not simply acting as an arbitrary basis function decomposition. To measure this, we exploited a key distinction between rotations and oscillations. Rotations require that each temporal basis function be embedded in a distinct dimension, so that each sine/cosine pair of temporal basis functions occupies a plane, and each different-frequency rotation occupies a different plane. In general, basis function expansions will not meet these requirements. We attempted therefore “recovered” the temporal basis functions from neural activity

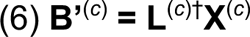

as where **B’**(c) is the recovered temporal basis functions for condition *c* and † indicates pseudo-inversion. We quantified the “recoverability” of temporal basis functions on condition *c* as the variance explained in the temporal basis functions by the projected neural activity. If this metric is 1, it suggests that the condition’s loading matrix is invertible, and therefore fulfills the requirements of rotations.

### Control for data smoothing

One potential concern is that the factorization described above works well on neural data simply because the data are smoothed before analysis, and smoothing limits the frequency spectrum and therefore allows neural activity to be reconstructed by a small number of temporal basis functions. To test these possibilities, for each trial of each condition independently we shuffled time bins before smoothing and averaging, to disrupt the temporal structure of motor cortex activity while preserving inter-neuron spike correlations and total spike counts. We then extracted temporal basis functions, with the number of temporal basis functions matched to the original dataset, and quantified the population variance explained and recoverability (defined above) of the temporal basis functions. We compared this null distribution to the data distribution using a Wilcoxon Signed Rank Test to test for differences in distributions.

### Control for stereotyped time course of behavior

One potential concern is that the factorization described above can find common frequencies between conditions simply because reaches take similar amounts of time, and motor cortex activity is likely modulated by movement onset, offset, and speed. If this hypothesis were correct, then time-warping reaches and the corresponding motor cortex activity to equalize reach duration should strengthen the recoverability of the temporal basis functions, as aspects of motor cortex activity are warped to become more aligned. To test this, we used linear time-warping to equalize all reaches at 600 ms duration, then time-warped motor cortex activity correspondingly. For the post-reach activity, in which motor cortex firing rates are still changing, we tested two methods: one warped the post-reach activity to uniformly last 250 ms, and the other truncated post-reach activity at 850 ms after movement onset. Both methods produced similar results; the latter variant is shown in Figure 3i. For both methods we then extracted the same number of temporal basis function as in the original dataset, and then quantified the recoverability of the temporal basis functions as a quantification of model fit. We then quantified the correlation between how much a reach condition’s duration differed from the mean and the recoverability. This was done for both the original and warped datasets.

### Subspace Excursion Angles

A set of vectors may be high-dimensional because they vary slightly from an otherwise low-dimensional plane, or because they are truly spread out and ‘fill’ the high-dimensional space. For example, 3-dimensional vectors can be chosen from within a cube or from within a narrow cone; either way, the space spanned is 3-dimensional, but if the vectors are chosen from the cube they will sweep ‘farther’ into the three dimensions. We attempted to distinguish these possibilities for the planes occupied by neural dynamics using a novel metric, Subspace Excursion Angles. More generally, in this analysis we sought to understand whether, for a set of low-dimensional subspaces {**S**^(1)^, …, **S**^(c)^} of a high-dimensional space, these subspaces varied only slightly relative to one another, forming small angles, or whether the objects truly “pivoted” far into many different dimensions and formed large angles with one another. Here, each **S**^(i)^ was the dimension(s) for a given component on a given condition (taken from the column(s) of loading matrices), embedded in high-dimensional neural state space. Existing methods of estimating dimensionality do not distinguish slight variations from more substantial variations, as long as the occupancy of each additional dimension is above the noise level. To better characterize the geometry of this set of subspaces, we took one of the observed subspaces **S**^(i)^ as a seed and found the greatest subspace angle (the principal angle) to any other observed subspace **S**^(j)^. We then replaced **S**^(i)^ with the union of **S**^(i)^ and the vector in **S**^(j)^ that formed the greatest angle with **S**^(i)^. We repeated this process to find the next greatest angle to the augmented **S**^(i)^, and so forth. This search was optimized over all possible sequences to find the sequence with the largest angles.

### Measuring the dimensionality of each rotation

We found that neural activity always contained rotations at a small number of fixed frequencies, but on different conditions neural dynamics potentially occupied dissimilar rotational planes. To determine whether each of the rotations occupied different planes on different conditions, or whether only some of the rotations did, we probed each rotation separately. To do so, we first grouped the pairs of identified temporal basis functions that shared the same frequency into rotations (a sine and cosine pair). Empirically, this produced 3-4 pairs of same-frequency sinusoidal temporal basis functions, and a single activity mode with zero frequency. We could correspondingly then group columns of each condition’s loading matrix that corresponded to single rotations. We then computed the alignment index for pairs of conditions for each rotation. For the state space location, the analysis was identical but with a single dimension instead of a pair. To determine whether one rotation ever used the same dimensions as a different frequency rotation on a different condition (Fig. 4b), we performed the same analysis but with the two planes coming from different rotations (different column pairs of **L**^(c)^). As before, the two planes came from different conditions. To put these alignment indices into context, it was necessary to understand what to expect due to estimation noise (gray distribution in Fig. 4b). To find the distribution expected from estimation noise, for each condition we randomly assigned trials into one of two partitions, then smoothed and trial-averaged each partition to estimate firing rates. We then used linear regression to identify the loading matrix for both partitions independently. Finally, we quantified the alignment index for rotational planes of the same frequency between these independent estimates of the same condition in either partition. We repeated this procedure 36 times to estimate a distribution of alignment indices.

### Explaining variability in jPCA-identified rotations

To link our findings with previous work, we applied jPCA to neural activity ^27^. As in the original work, we applied jPCA with mean-subtraction, a soft-normalization constant of 5, and “PC smoothing” (which we note is a single-neuron version of our same matrix decomposition). Based on visual inspection and the results in the original work, we kept the first two of the discovered jPCA planes for M1 as “rotational”, while for PMd we kept only the first plane. To quantify the fit of jPCA to the total activity, we computed the percentage of population variance contained in the plane(s). To quantify total rotational activity on condition *c* we quantified the variance due to rotational temporal basis functions from the decomposition in our equation 3. To compare to jPCA, prior to quantifying the variance or extracting temporal basis functions we soft-normalized and mean subtracted activity identically to jPCA. We then quantified the Pearson correlation of this variance measure with the variance in each jPCA plane across conditions, for each retained jPCA plane independently. To quantify the alignment between neural activity and the jPCA plane on condition *c* (Fig. 4f), we computed the alignment index between the total rotations in neural activity and the jPCA plane for each condition. We then quantified the Pearson correlation of the alignment index with the variance in each jPCA plane across conditions, for each retained jPCA plane independently.

### Predicting rotations from other rotations or the state space location

On a single condition, that condition’s loading matrix describes the state space location and each rotation’s parameters: the rotational plane, along with the initial location of the population state within the plane. To demonstrate the rotations’ parameters were not independent, we predicted the parameters of one rotation from another’s on the same condition. Each rotation’s parameters on a single condition are described by a pair of columns in the loading matrix. We therefore used linear regression to predict the pair of columns describing one rotation’s parameters from the pair of columns describing a different rotation’s parameters. This was done by vectorizing each pair of columns for the predictor rotation and the predicted rotation, with each condition acting as an observation, then using ridge regression (lambda = 0.1) to relate the two rotations. We cross-validated our model with 6-fold cross validation: fitting temporal basis functions, loading matrices, and the regression model to 5/6 of the conditions, then testing on the remaining 1/6. To quantify model performance, we reshaped the output of the model back into a pair of column vectors, and then multiplied by the two associated temporal basis functions to produce the “predicted rotation”. We correspondingly multiplied the two empirical columns of the loading matrix by the same temporal basis functions to produce the “true rotation”. We then quantified the variance explained in the true rotation by the predicted rotation. We repeated this procedure over all conditions and pairs of rotations.

We similarly fit a nonlinear version of this model using kernel ridge regression ^57^ (lambda = 0.1, Gaussian Kernel, length-scale = 400). Parameters for this and all other kernel regressions were optimized by grid search. As a null distribution, we independently shuffled predictor and predicted rotations between conditions, quantified variance explained by model output, and tested for difference in the true distributions used a Wilcoxon Signed Rank Test to test for differences in distributions.

To extend this procedure to the state space location, we used the column of the loading matrix associated with state space location as a predictor, and predicted the (vectorized) remainder of the loading matrix using ridge regression (lambda = 0.1). We identically used 6-fold cross validation for this model. To quantify model performance, we concatenated the observed state space location column with the (reshaped) output of the model, multiplied the resulting matrix by the temporal basis functions, then computed variance explained in that condition’s neural activity. We similarly fit a nonlinear version of this model using kernel ridge regression (lambda = 0.1, Gaussian Kernel, length-scale = 400). As a null distribution, we independently shuffled state space locations and rotations between conditions, quantified variance explained by model output, and tested for difference in the true distributions used a Wilcoxon Signed Rank Test to test for differences in distributions.

### LDR Encoding

On a single condition, the state space location and rotation’s parameters are given by that condition’s loading matrix. By contrast, the temporal basis functions describing the frequencies of these rotations are common across conditions. To predict neural activity from kinematics, we therefore used linear regression to predict a condition’s loading matrix from that condition’s kinematics. To both regularize the model and to allow for model interpretability, we reduced the dimensionality of the kinematics on each condition using linear dimensionality reduction. We regressed each condition’s hand position over time against two kinematic basis functions to extract four “reach parameters” that described target location and reach curvature along the x- and y-axis. We vectorized each condition’s loading matrix, and used ridge regression (lambda = 0.1) to fit a map from these four coefficients to a condition’s loading matrix. We used 6-fold cross-validation, fitting temporal basis functions, loading matrices, and the regression model to 5/6 of the conditions, then quantifying model performance on the remaining 1/6 of the conditions. We quantified model performance by reshaping model output, multiplying by the temporal basis functions, then quantifying variance explained in neural activity. We similarly fit a nonlinear version of this model using kernel regression (Ornstein-Uhlenbeck kernel, length-scale = 10^3^). As a null distribution, we shuffled neural activity and kinematics between conditions, quantified variance explained by model output, and tested for difference from the true distributions using a Wilcoxon Signed Rank Test.

### Extracting kinematic basis functions

To perform sequence-to-sequence mapping of an entire neural trajectory to an entire kinematic trajectory, we required a low-dimensional and meaningful summary representation of reaching kinematics over the entire trial. We therefore performed an SVD on the x- and y-coordinate velocity of the hand on all conditions in all our datasets (Fig. 6d). The top two velocity singular vectors were interpretable. The first velocity singular vector was a unimodal trace, with rapid rise from zero beginning around movement onset, peaking around 250 ms after movement onset, and then slower decay toward zero. The second velocity singular vector was a bi-phasic trace, with alternating up and down modes, and approximately represented the derivative of the first vector. The first velocity singular vector therefore mostly contained information about the straight reaching velocity component, while the second velocity singular vector contained information about reach curvature or target overshoot. We fit these two kinematic basis functions as a skew-normal curve and the derivative of a skew-normal curve. Decoding results were similar if the raw vectors from SVD were used, but this fit ensures that position and velocity variants of the model are identical. We optimized over parameters using gradient descent. After fitting these components, we analytically integrated them to obtain the equivalent basis functions over position.

### Controlling for nonlinear LDR encoding

One potential concern in nonlinear regression models is that the model will collapse points in the test set towards the nearest points in the training set, acting not as a continuous regression model but as a nearest-neighbor regression model. To control for this possibility, we divided target locations into sextants. We trained nonlinear LDR encoding models on 5/6 of the sextants and tested on the held-out sextant. For comparison, we trained a nonlinear LDR encoding model that used nearest neighbor regression in place of kernel regression with the same cross-validation. This explicitly compares nonlinear regression’s performance to memorization of the training set. We compared distributions of variance explained using either approach with a Wilcoxon Signed Rank Test.

### Comparing the LDR encoding with standard instantaneous encoding models

To compare our encoding model based on LDR to standard, instantaneous tuning models, we fit several forms of encoding model to motor cortex activity. In particular, we compared:

- Linear regression to predict motor cortex activity as a linear combination of baseline firing rate, position, extent, velocity, speed, velocity angle, and acceleration.
- Kernel regression to predict motor cortex activity from hand position.
- Kernel regression to predict motor cortex activity from hand velocity.

For all regressions, we included a 100 ms causal lag between kinematics and neural activity. For linear regression, we used ridge regression (lambda = 0.1). For kernel regression, we used the Ornstein-Uhlenbeck kernel with a length-scale of 10^12^ for position encoding and 10^11^ for velocity encoding. For each encoding method, we then quantified the variance explained for motor cortex activity, using each condition as a data point for the distributions. Finally, we compared these distributions to the distributions obtained from LDR encoding using a Wilcoxon Signed Rank Test.

### LDR-based decoding

We fit an LDR-based linear decoder that directly used the state space location and rotation orientations to decode reach kinematics. The predictors for this decoder consisted of the vectorized loading matrices of each condition. The targets of the model consisted of the four reach parameters described earlier, which capture target position and reach curvature. We used 6-fold cross-validation: fitting temporal basis functions, loading matrices, and the decoder to 1/2 of the conditions (trial-averaged). For the remaining 1/2 of the conditions, we decoded from single-trials rather than trial-averaged activity. To decode, we estimated the loading matrix for each trial by pseudoinverting the temporal basis functions and multiplying with the trial’s spike counts (see Supplement). We then vectorized this estimated loading matrix and used the model to predict the four coefficients. Finally, we multiplied these coefficients by the kinematic basis functions to produce an estimate of reach position over time. We quantified model fit as the variance explained in reach position over time. As a null distribution, we shuffled neural activity and kinematics between conditions, quantified variance explained, and compared with the true distributions using a Wilcoxon Signed Rank Test.

### Comparing LDR-based decoding to instantaneous decoding methods

We benchmarked our decoding methods against advanced versions of more traditional decoding models, which decode the value of kinematic parameters at an instant in time from motor cortex firing rates at an earlier instant in time. As exemplary instantaneous decoding methods, we used:

- Linear regression to decode hand position or hand velocity, with a lag of 100 ms, on spiking activity smoothed with a 20 or 100 millisecond Gaussian kernel.
- Kernel regression to decode hand position or hand velocity, with a lag of 100 ms, on spiking activity smoothed with a 20 or 100 millisecond Gaussian kernel, trained on a subset of single-trial data.
- Same as above, but trained on trial-averaged activity.
- Linear or kernel regression to decode hand position or hand velocity, with a lag of 100 ms, on single-trial trajectories inferred by Gaussian Process Factor Analysis (GPFA).

For linear regression, we used ridge regression (lambda = 1). For kernel regression, we used the Ornstein-Uhlenbeck kernel with a length-scale of 10^2^ and an L2-penalty of 0.1. For each decoding method, we quantified the variance explained in hand position for each condition. We compared each of these distributions to the data distribution using a Wilcoxon Signed Rank Test.

### Quantifying the trial-to-trial fidelity of decoding within condition

An ideal kinematic decoder would follow variations in individual movements, not categorically decode condition identity and output kinematics corresponding to the condition. We wished to test whether the LDR-based decoder acted akin to a nearest neighbor classifier, detecting which condition a trial was most similar to and returning the corresponding average reach. To test whether our decoder was continuous or categorical, we asked whether it could decode the residuals of the reach parameters, within a condition. Specifically, we considered the third and fourth reach parameters, which describe curvature. We then quantified the Pearson correlations between the decoded and true reach parameters.

## Supplement

### Basis function representation of a diagonalizable linear dynamical system

Linear dynamical systems are defined by a transfer matrix, **M**, that maps the state of the dynamical forward in time. This can be defined in continuous time as

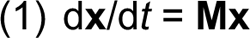

so that the derivative of the state is just a linear transformation on the current state. From some initial state **x**0, the state at time *t* can be expressed as

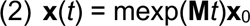

where mexp() refers to the matrix exponential. In this section, we will work in continuous time, though our argument generalizes to discrete time naturally. Under a few assumptions, the transfer matrix **M** can be diagonalized

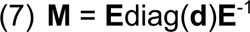

where **E** is the eigenvectors of the dynamics, **d** is a vector of the eigenvalues, and diag() returns a diagonal matrix with the argument along the diagonal. If **M** is diagonalizable, then the state at time *t* can be expressed more simply as

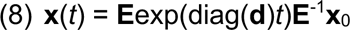

where exp() is the exponential function applied to each element in its argument. Define the vector

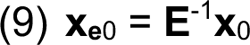

as the initial state in the eigenvector basis. Using that, for two vectors **z** and **y** of identical dimensions,

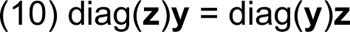

then (8) can be re-written as

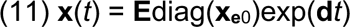

so that exp(**d***t*) is now a column-vector. By further defining the terms

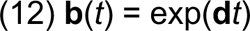

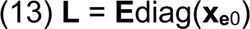

(11) can be re-written as

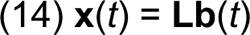

so that the state at time *t* can be expressed as the product of loading matrix and a set of temporal basis functions. This factorization isolates the effect of the eigenvalues in determining the “temporal” characteristics of the population into **b**. **L**, by contrast, expresses both the effect of the eigenvectors of determining “where” in state space dynamics occur, and the initial state in setting the phases and magnitudes of the dynamics.

### Fitting motor cortex activity with conserved rotations, but varying rotational planes

In this study, we required a method to identify a model of motor cortex activity where rotations in motor cortex activity were conserved in their temporal characteristics (identical eigenvalues), but occupied different planes on different conditions (different eigenvectors). In this model, motor cortex activity on single conditions is an LDS. This system is, however, nonlinear across conditions, thus disallowing simply regressing the state’s derivatives against the state. We exploit the basis-function representation of LDSs to work around this issue. As conditions share eigenvalues, they share temporal basis functions. This means that two conditions, *i* and *j*, have states as a function of time *t*

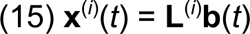

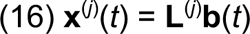

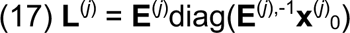

where **x**^(*j*)^(*t*) and **x**^(*i*)^(*t*) is the firing rates at time *t* on conditions *j* and *i* respectively, **E**^(*j*)^ is the matrix of eigenvectors of the transfer matrix for condition *j*, and **x**^(*j*)^0 is the initial state on condition *j*. Usefully, while the loading matrices in (15) and (16) differ, the temporal basis functions are conserved between conditions because the eigenvalues are conserved. This means that activity on each condition is just a different linear transformation on the same temporal basis functions. Fitting this model therefore reduces to identifying an optimal set of temporal basis functions that explain neural activity. These can be obtained by the temporal singular vectors of the neural activity concatenated into a (*neurons* x *conditions*)-by-*time* matrix, or equivalently the eigenvectors of the Gramian matrix over time. This is comparable to similar operations over neurons that are used to extract the dimensions of highest variance in neural state space.

---

**Extended Data Figure 1:**
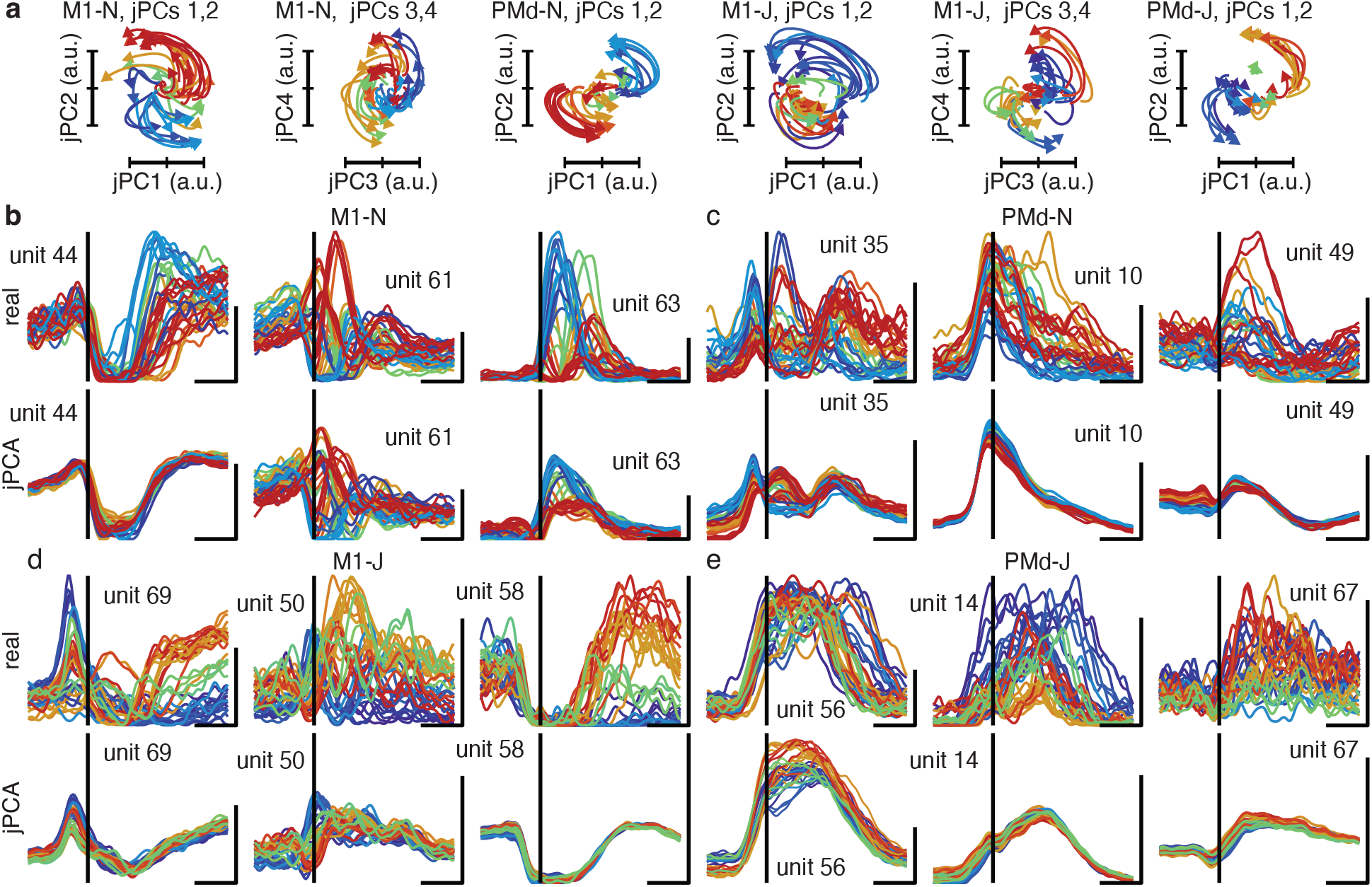
jPCA rotations incompletely describe motor cortex activity. **a**, Rotations uncovered by jPCA across datasets. jPCA identified two rotational planes in M1 activity, and one rotational plane in PMd. **b**, Multi-unit PSTHs and reconstruction by jPCA rotations and mean activity across conditions. Rotations captured little of the multi-phasic modulation of neural activity during reaching. Traces colored by target angle.

**Extended Data Figure 2:**
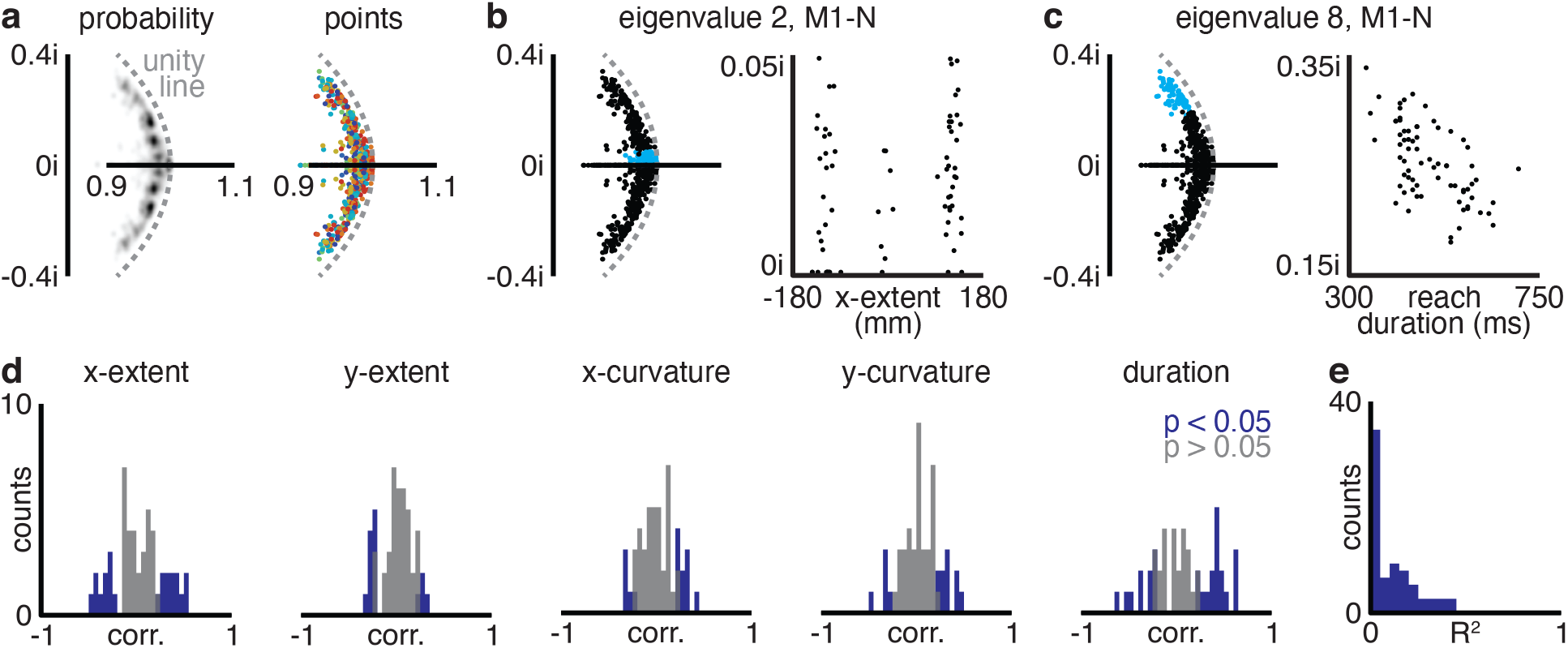
Eigenvalues of motor cortical dynamics correlate modestly with reach parameters. **a**, Probability distribution of eigenvalues for single-condition dynamics (*left*) and eigenvalues colored by target angle (*right*). **b**, Second eigenvalues of single-condition dynamics (blue, *left*), and x-extent of reach vs. the imaginary component of the second eigenvalue (*right*). No obvious relationship is present. **c**, Eighth eigenvalues (corresponding to the highest-frequencies that were reliable) of single-condition dynamics (blue, *left*), and reach duration vs. the imaginary component of the eighth eigenvalue (*right*). Data in **a**-**c** taken from M1-N. **d**, Histogram of correlation coefficients of reach parameters with each real and imaginary component of each eigenvalue. **e**, Histogram of R^2^ for linear models predicting real and imaginary components of eigenvalues from reach parameters. Data in **d,e** pooled across datasets.

**Extended Data Figure 3:**
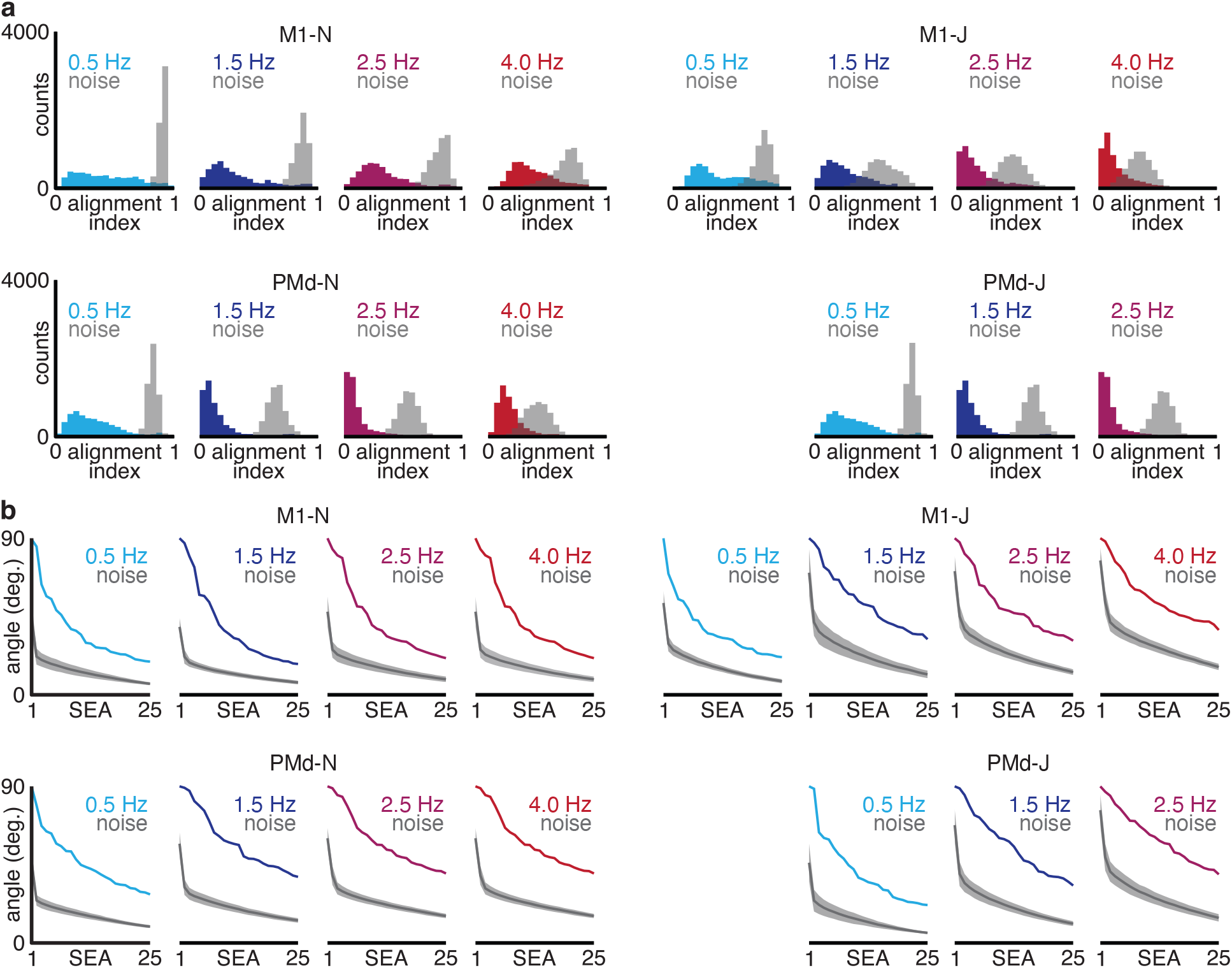
Rotational planes differ between conditions. **a**, Alignment indices between corresponding (same-frequency) rotational planes on pairs of conditions (*colored*) and distribution expected due to estimation noise (*gray*). **b**, Subspace excursion angles for corresponding rotational planes across conditions (*colored*), along with angles expected due to estimation noise (*gray*; Line, mean; shaded, 1 standard deviation).

**Extended Data Figure 4:**
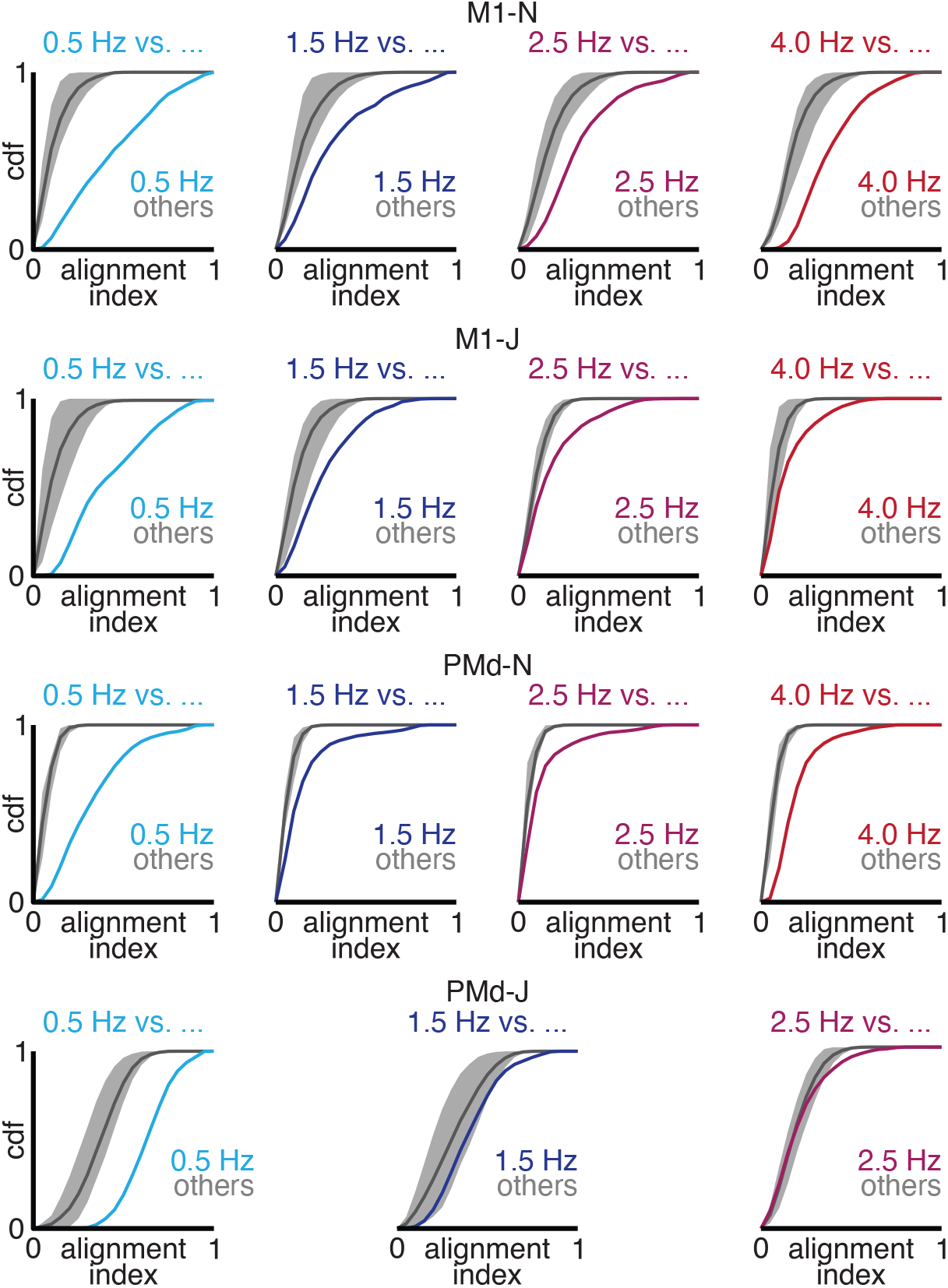
Rotations occupied distinct subspaces in neural state space. Cumulative distribution function of alignment index between different-frequency rotational planes (*gray*) and same-frequency rotational planes (*colored*) on pairs of conditions. Same-frequency rotational planes were more aligned than different-frequency rotational planes.

**Extended Data Figure 5:**
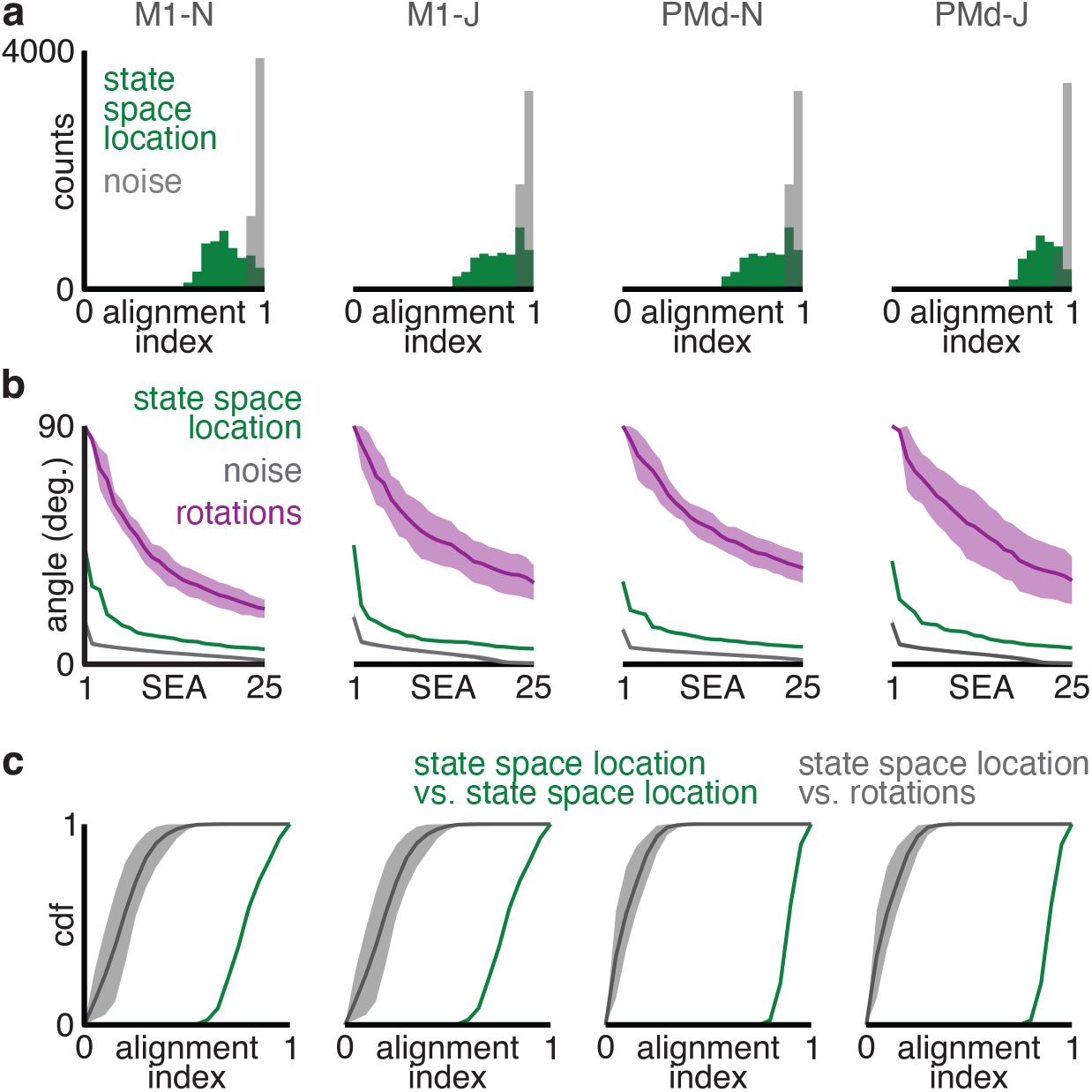
State space location differed between conditions. **a**, Alignment indices between state space locations on pairs of conditions (*green*) and distribution expected due to estimation noise (*gray*). **b**, Subspace excursion angles for state space locations across conditions (*green*), along with angles expected by estimation noise (*gray*) and angles formed by rotations (*purple*). Line, mean; shaded, 1 standard deviation. **c**, Cumulative distribution function of alignment index between rotations and state space locations (*gray*) and state space locations compared to state space locations (*green*) on pairs of conditions. State space locations were more aligned with themselves than with rotations.

**Extended Data Figure 6:**
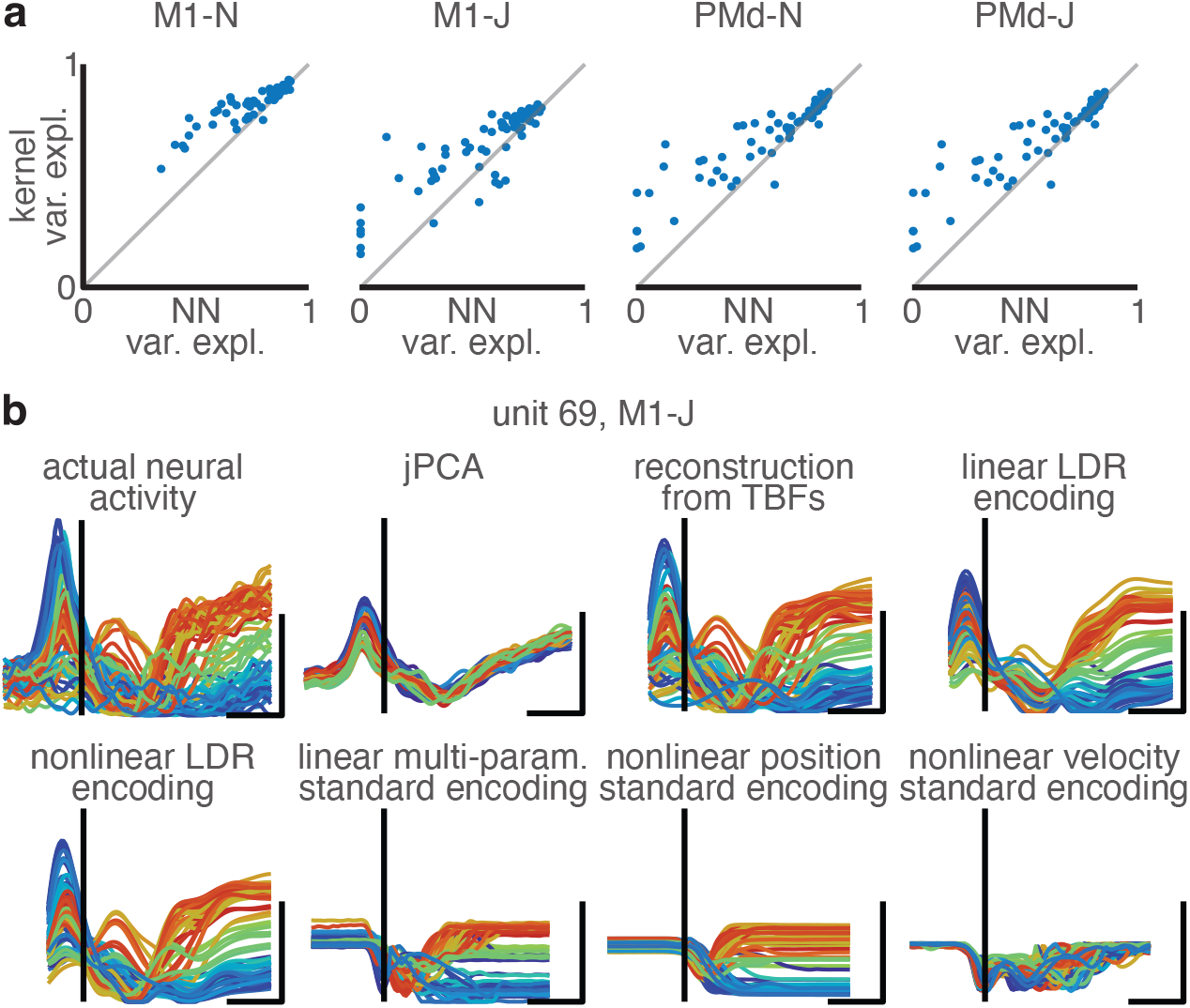
Reach kinematics are sufficient to predict motor cortex dynamics. **a**, To demonstrate that LDR encoding used continuous features to determine test condition dynamics, rather than simply memorizing the training set, we compared kernel regression (a continuous model) with nearest-neighbor regression (which memorizes the training set). Conditions were sorted by target angle. Angles were divided into sextants for cross-validation. The higher performance of kernel regression indicates that nonlinear LDR encoding was not merely memorization of the training set. **b**, Example unit for the 7 models of neural activity fit in this study.

